# *Drosophila Bchs* overexpression recapitulates human *WDFY3* neurodevelopmental phenotypes with implications for glial cell involvement in altered head circumference

**DOI:** 10.1101/2023.10.16.562295

**Authors:** Marek B. Körner, Akhil Velluva, Linnaeus Bundalian, Knut Krohn, Kathleen Schön, Isabell Schumann, Jessica Kromp, Andreas S. Thum, Antje Garten, Julia Hentschel, Rami Abou Jamra, Achmed Mrestani, Nicole Scholz, Tobias Langenhan, Diana Le Duc

## Abstract

The autophagy adaptor *WDFY3* is linked to neurodevelopmental delay and altered brain size. Loss-of-function variants are associated with an increased brain size in both humans and mice. We thus, hypothesized that the microcephaly observed in some of the patients may be related to a gain-of-function of the *WDFY3* gene product. While the role of *WDFY3* loss-of-function has been studied extensively in neurons, little is known about the effects of *WDFY3* overexpression in different neural cell types. We utilized a *Drosophila melanogaster* overexpression model to investigate the effect of the *WDFY3* ortholog *Bchs* (*blue cheese*) on development, CNS size, and gene expression profiles. Glial and neuronal overexpression of *Bchs* impaired CNS development, locomotion and autophagy. Glial overexpression of *Bchs* also altered CNS size significantly. We identified 79 genes that were differentially expressed and overlapped in flies that overexpress *Bchs* in glial and neuronal cells, respectively. Additionally, upon neuronal *Bchs* overexpression differentially expressed genes clustered in gene ontology categories associated with autophagy and mitochondria. Our data indicate that *WDFY3*/*Bchs* overexpression in both neurons and glial cells results in impaired neural development, which corresponds to symptoms observed in *WDFY3*-related neurodevelopmental delay.

## Introduction

Over the past decade, a growing body of evidence, including functional analyses and patient related data, has provided substantial support for the involvement of *WDFY3* in neurodevelopmental disorders (Le Duc *et al*. 2019; Stessman *et al*. 2017; Wang *et al*. 2016). *WDFY3* encodes an autophagosomal scaffolding protein involved in targeted recruitment and destruction of macromolecular components including aggregation-prone proteins (Clausen *et al*. 2010; Filimonenko *et al*. 2010; Finley *et al*. 2003; Simonsen *et al*. 2004).

Previous studies in mice demonstrated that *Wdfy3* regulates neurodevelopmental processes such as neuronal connectivity, proliferation, migration, and synaptic morphology (Dragich *et al*. 2016; Orosco *et al*. 2014; Schaaf *et al*. 2022; Søreng *et al*. 2022). Loss-of-function variants of *WDFY3* or its *Drosophila* ortholog *blue cheese* (*Bchs*) result in neurodegeneration and protein aggregation, indicating autophagic defects (Clausen *et al*. 2010; Filimonenko *et al*. 2010; Finley *et al*. 2003; Fox *et al*. 2020; Han *et al*. 2015; Hebbar *et al*. 2015; Lim & Kraut 2009; Simonsen *et al*. 2004). The increased head circumference, but also a decreased learning capacity observed in affected human individuals, were faithfully recapitulated in heterozygotes of a *Wdfy3* knockout mouse model, while homozygotes of hypomorphic *Wdfy3* alleles showed perinatal lethality (Dragich *et al*. 2016; Le Duc *et al*. 2019; Orosco *et al*. 2014). However, while loss-of-function variants were associated with increased head circumference, at least two variants have been identified in individuals with microcephaly (Kadir *et al*. 2016; Le Duc *et al*. 2019). We, thus, hypothesized that *WDFY3* loss– and gain-of-function genotypes may result in opposing phenotypes in respect to brain size.

Currently an increasing number of studies concentrate on loss of *WDFY3*/*Bchs* and its effects on the nervous system with a focus on neuronal impairment. Although glial cells are known to play an essential role for neuronal function and neurodevelopment (Bittern *et al*. 2021; Kim *et al*. 2020; Lago-Baldaia *et al*. 2020; Rahman *et al*. 2022), they have received rather little attention in the effort to unravel *WDFY3*-associated pathomechanisms. So far, it was shown that loss of *Wdfy3* is accompanied with mislocalisation of glial guidepost cells, which provide guidance cues for the formation of axonal tracts (Dragich *et al*. 2016). Hypomorphic *Wdfy3* alleles also increase symmetric proliferative divisions of radial glial cells, neural stem cells which give rise to neurons and glia (Orosco *et al*. 2014). Further, *Wdfy3* is involved in the turnover of oligodendrocytic myelin sheaths (Aber *et al*. 2022). Single-cell RNA-sequencing (RNA-seq) demonstrated an approximately 10× higher *WDFY3* expression in neurons and glial cells compared to all other cells (nTPM > 200) with oligodendrocytes showing the highest expression (nTPM > 400) (Karlsson *et al*. 2021). Hence, glial cells may play an important role in the pathophysiology of *WDFY3*-related neurodevelopmental disorders.

To address the unknown role of glial cells in *WDFY3*-related pathologies and to understand whether *WDFY3* overexpression as proxy for a *WDFY3* gain-of-function condition may be related to neurodevelopmental disorders, here we investigated the effects of *Bchs* overexpression in glial cells and neurons. While both glial and neuronal *Bchs* overexpression impaired neural development, locomotion, and autophagy, central nervous system (CNS) size was altered only after overexpression in glial cells. Further, based on transcriptomics analyses we identified differentially expressed genes in glial and neuronal *Bchs* overexpression flies compared to the respective controls. We found an overlap of 79 differentially expressed genes in both *Bchs* overexpression conditions, which may be involved in the pathological mechanism that unfolds upon *WDFY3* dysfunction.

## Results

### Developmental delay in glial Bchs overexpression flies

A common symptom of probands carrying a pathogenic *WDFY3* variant is neurodevelopmental delay (Le Duc *et al*. 2019; Stessman *et al*. 2017; Wang *et al*. 2016). In flies, both loss-of-function and overexpression of the *WDFY3* ortholog *Bchs* were previously shown to impair neuronal function (Finley *et al*. 2003; Hebbar *et al*. 2015; Khodosh *et al*. 2006; Kraut *et al*. 2001; Kriston-Vizi *et al*. 2011; Lim & Kraut 2009; Sim *et al*. 2019; Stessman *et al*. 2017). However, not much is known about how *Bchs* dysregulation impacts glial cells. To better understand the relevance of *Bchs* in different neural cell types, we tested the effect of *Bchs* overexpression, as a proxy for gain-of-function effects, in glial cells (*repo-Gal4*) and neurons (*nSyb-Gal4*) on development, locomotion, CNS morphology, and autophagy (DiAntonio *et al*. 1993; Sepp *et al*. 2001). We found that panglial *Bchs* overexpression delayed the development from egg to adult fly (Fig 1A). Importantly, Mendelian ratios of the adult F1 generation of crossing *Gal4/Sb* with *UAS-bchs::HA/Sb* for panglial and panneuronal *Bchs* overexpression corresponded to the expected ratios, but ubiquitous *Bchs* overexpression (*act5C-Gal4*) was lethal (Fig S1). The developmental time of neuronal *Bchs* overexpression animals was further investigated to exclude a delay in early developmental steps, which cannot be detected at the stage of adult eclosion. The timepoint of larval hatching was not delayed in animals overexpressing *Bchs* in neurons, indicating normal embryonal development (Fig S2A). However, a significantly reduced size of neuronal *Bchs*-overexpressing larvae was observed in early larval development (62.44 h and 100.8 h after egg laying), but not in later larval stages (153 h after egg laying) (Fig S2B). These data suggest that neuronal *Bchs* overexpression causes changes in development during early larval stages.

**Fig 1.**
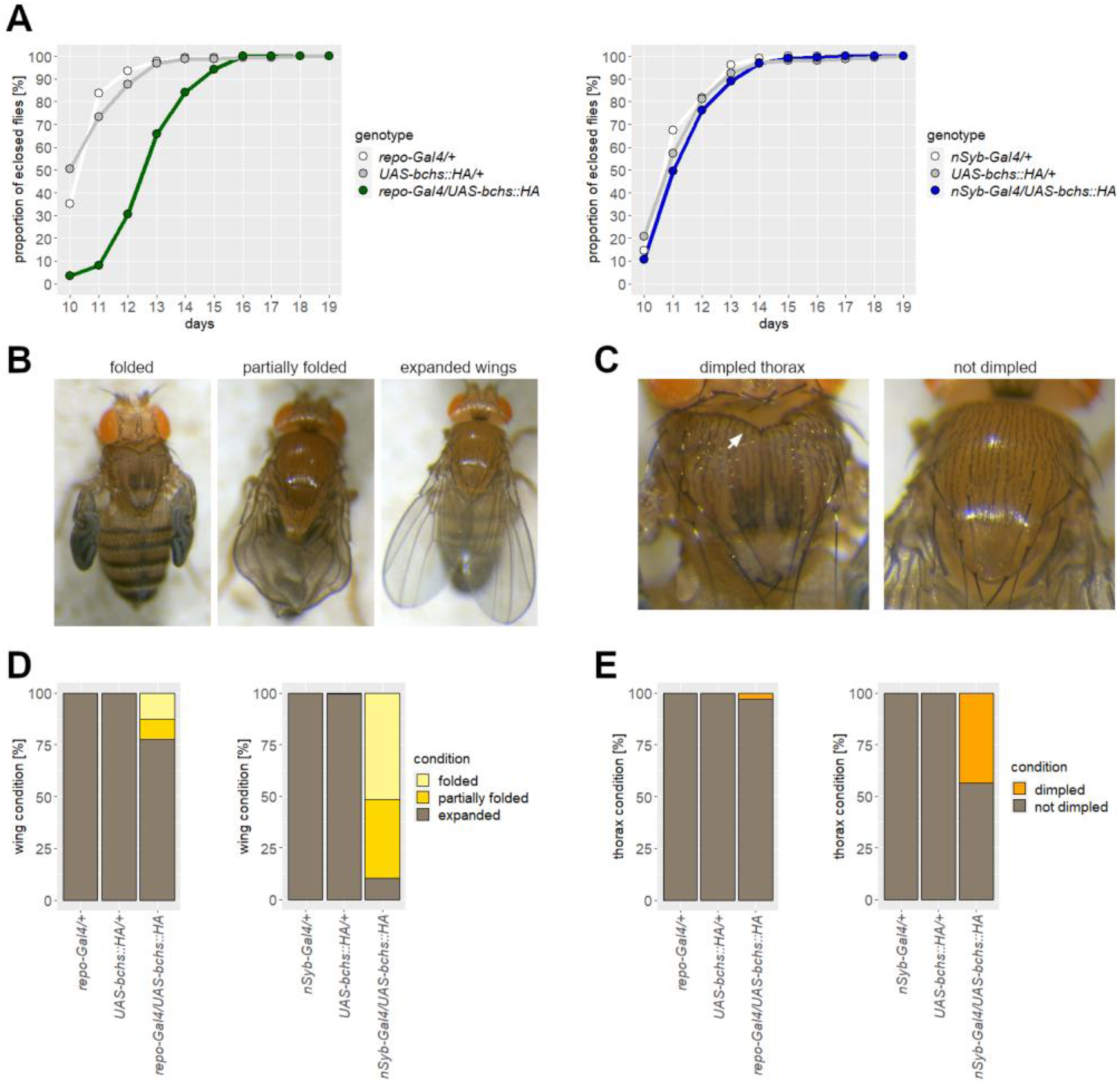
Developmental deficits in Bchs overexpression flies. A) Glial *Bchs* overexpression (left, *repo-Gal4/UAS-bchs::HA*, green) prolonged the developmental time from egg to eclosion of adult flies, unlike neuronal overexpression (right, *nSyb-Gal4/UAS-bchs::HA*, blue), in comparison to controls (left: *repo-Gal4/+* white, *UAS-bchs::HA/+* gray) (right: *nSyb-Gal4/+* white, *UAS-bchs::HA/+* gray). Flies of 5 vials were pooled. n > 88. B, C) Images of flies overexpressing *Bchs* in neurons. *Bchs* overexpression caused wing expansion and dimpled thorax defects (arrow). D, E) Quantification of wing (D) and thorax (E) defects. Both defects had a higher rate in neuronal than in glial *Bchs* overexpression. n > 72.

Panneuronal *Bchs* overexpression caused deficits in the development of the wings and thorax (Fig 1B,E). Adult flies did not properly expand their wings (∼90 %) and displayed a dimpled thorax (∼43 %). In contrast, only a small percentage of flies overexpressing *Bchs* in glial cells (wing: ∼22 %, thorax: 3 %) presented with those deficits (Fig 1D,E). Crustacean cardioactive peptide (CCAP) neurons are known to play an essential role in wing expansion (Luan *et al*. 2006; Park *et al*. 2003). Therefore, we hypothesized that *Bchs* overexpression in this subtype of neurons caused the wing and thorax abnormalities. Driving *Bchs* overexpression specifically only in CCAP neurons (*CCAP-Gal4*) resulted in almost complete penetrance of those defects (wing: ∼100 %, thorax: ∼99 %), as opposed to *Bchs* overexpression in motoneurons (*ok6-Gal4*, wing: ∼19 %, thorax: ∼1 %)) (Fig S1). Hence, CCAP neurons are sensitive to *Bchs* overexpression.

Since probands with *WDFY3* variants show impaired motor coordination (Le Duc *et al*. 2019), we also tested larval locomotion in our fly models. Both panglial and panneuronal *Bchs* overexpression decreased larval crawling velocities indicating a locomotion deficit (Fig 2C,F). Overexpressing *Bchs* only in the subset of motoneurons (*ok6-Gal4*) was sufficient to slow larval movement (Fig S4). Together these data show that glial and neuronal *Bchs* overexpression impacts fly development and locomotion.

**Fig 2.**
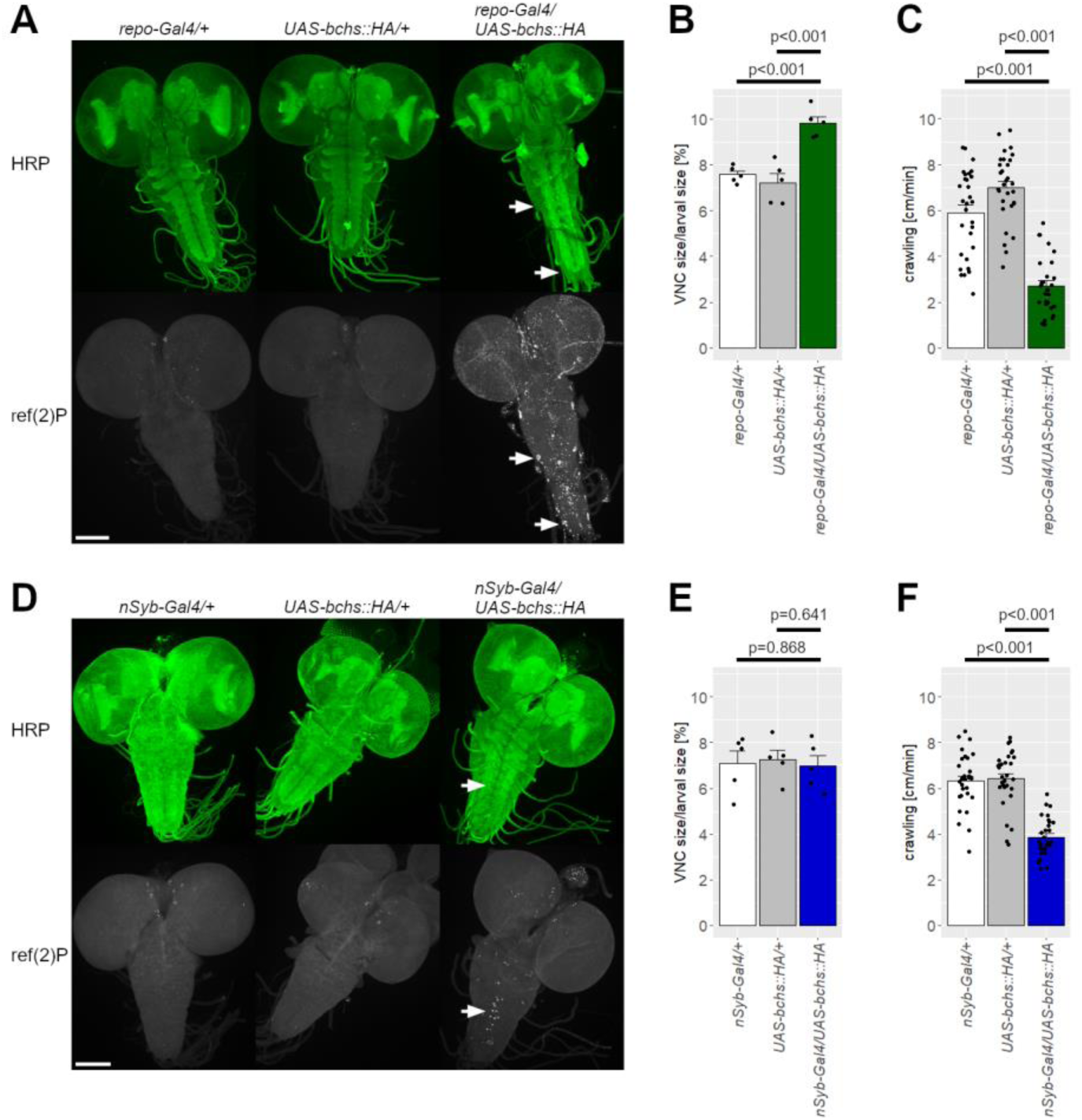
Larval locomotion and VNC length were affected by glial Bchs overexpression. A, D) Larval brains were stained against HRP (top) and ref(2)P (bottom). *Bchs* overexpression in glial cells (A, *repo-Gal4/UAS-bchs::HA,* right) and neurons (D, *nSyb-Gal4/UAS-bchs::HA,* right) caused accumulation of ref(2)P, in contrast to controls. Arrows exemplary point out ref(2)P aggregates. Scale bar: 100 µm. B, E) Quantification of VNC length normalized to larval length. Increased VNC length was observed in glial *Bchs* overexpression (B, green) larvae but not neuronal overexpression (E, blue). n =5. C, F) Overexpression of *Bchs* in glial cells (C, green) or neurons (F, blue) reduced the crawling velocity. n > 29. B, C, E, F) Data are shown as mean ± SEM.

### Glial Bchs regulates CNS size and glial cell number

Another symptom in carriers of pathogenic *WDFY3* variants is their abnormal head circumference (Le Duc *et al*. 2019; Stessman *et al*. 2017; Wang *et al*. 2016). It was previously described that the *Bchs* loss-of-function mutant *bchs^58^* presents with decreased larval and adult brain volumes, while neuronal *Bchs* overexpression by *elav-Gal4* causes an increased larval brain volume (Finley *et al*. 2003; Kriston-Vizi *et al*. 2011). Further, fly pharates ubiquitously overexpressing human *WDFY3* carrying a missense variant that is linked to microcephaly in human individuals, displayed smaller brain volumes (Kadir *et al*. 2016).

In our setup, neuronal *Bchs* overexpression did not alter the brain size to fly length ratio in adult flies or the ventral nerve cord (VNC) length in larvae and adults (Fig 2E,3E,F). However, we found that adult flies overexpressing *Bchs* in glial cells had a decreased brain size to fly length ratio (Fig 3B), while neither brain size nor fly length alone were significantly altered in comparison to controls (Fig S5A,B). Further, glial *Bchs* overexpression markedly increased the longitudinal VNC length in larvae and adult flies (Fig 2B,3C). In adults, the length of the abdominal but not thoracic neuromeres of the VNC was elongated (Fig S5C). We then sought to identify the glial cell type responsible for VNC elongation by enforcing *Bchs* overexpression in glial subtypes through specific *Gal4* drivers. We found that *Bchs* overexpression in subperineural glial cells (*rL82-Gal4*) was sufficient to increase the larval VNC length, while overexpression in perineural (*c527-Gal4*) or wrapping glia (*nrv2-Gal4*) was not (Fig S6). Subperineural glial cells are essential for the blood brain barrier (BBB) and form septate junctions (Baumgartner *et al*. 1996; Stork *et al*. 2008). These data demonstrate that glial *Bchs* plays a role in regulating CNS size.

**Fig 3.**
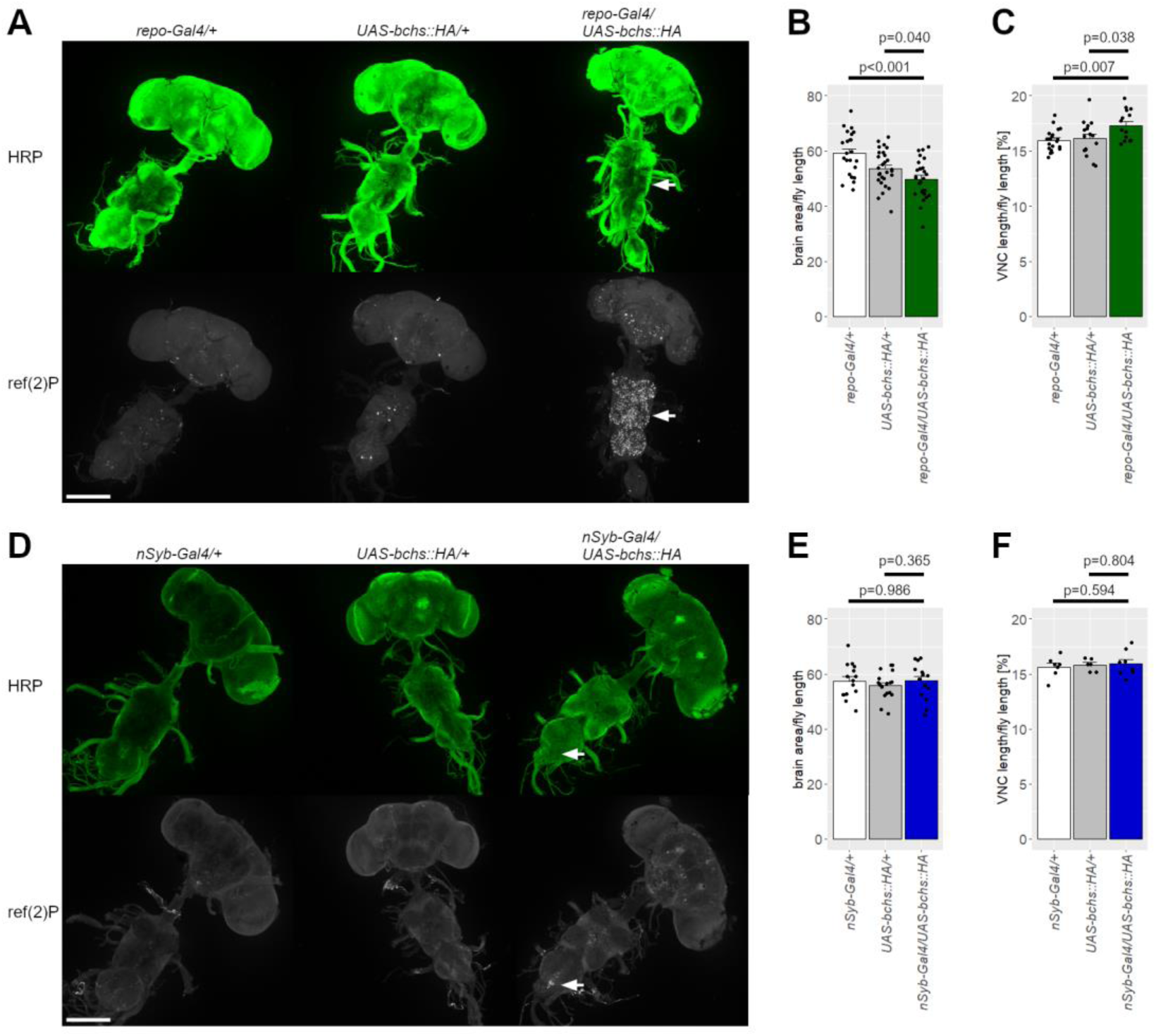
Glial Bchs overexpression affected adult CNS size. A, D) Adult CNSs were stained against HRP (top) and ref(2)P (bottom). *Bchs* overexpression in glial cells (A, *repo-Gal4/UAS-bchs::HA,* right) caused accumulation of ref(2)P in the brain and thoracic neuromeres of the VNC. Neuronal *Bchs* overexpression (D, *nSyb-Gal4/UAS-bchs::HA,* right) led to ref(2)P accumulation in the brain and posterior region of the VNC. Arrows indicate ref(2)P accumulation. Scale bar: 200 µm. B, E) Quantification of brain size normalized to fly length. A decreased brain size to fly length ratio was noted in glial *Bchs* overexpression adults (B, green) but not for neuronal overexpression (E, blue). B) n ≥ 23. E) n ≥ 15. C, F) Overexpression of *Bchs* in glial cells (C, green), but not in neurons (F, blue), elongated the VNC length (normalized to fly length). C) n ≥ 15. F) n ≥ 6. B, C, E, F) Data are shown as mean ± SEM.

Elongation of the VNC may be caused by an increased number of glial cells. Therefore, we quantified the number of glial cells in the larval brain using an anti-repo antibody, which is a panglial cell marker (Xiong *et al*. 1994). Panglial *Bchs* overexpression significantly increased the number of repo^+^ nuclei but not their density (Fig 4A,B). We additionally observed an increased number of glial cells in the peripheral nervous system (Fig S7). In conclusion, *Bchs* overexpression can act on glial proliferation or apoptosis inside and outside the CNS.

**Fig 4.**
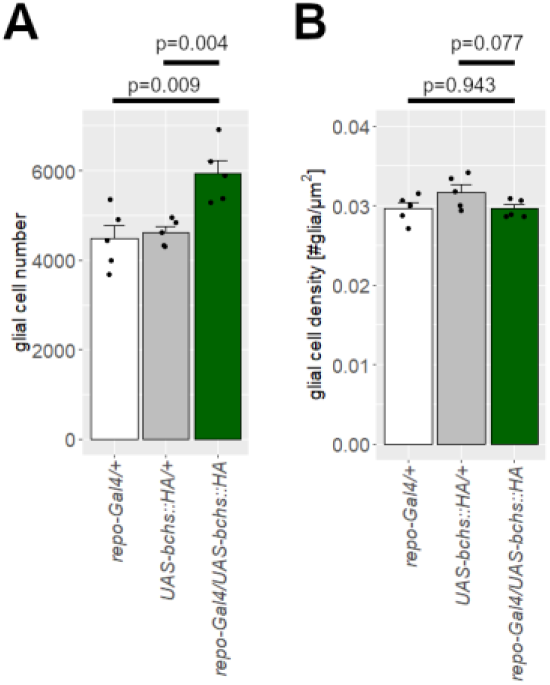
Glial cell number in larval CNS increased by glial Bchs overexpression. A) Glial nuclei in the larval CNS were counted by staining against repo. B) Glial cell density was determined by dividing the glial cell number by the larval CNS area. A, B) Glial *Bchs* overexpression raised the glial cell number but did not alter the glial cell density. n = 5. Data are shown as mean ± SEM.

### Bchs overexpression causes protein accumulation and a shift towards non-acidic autophagic vesicles

The autophagy adaptor ref(2)P links ubiquitinated proteins to autophagosomes via interactions with Atg8a, which is anchored to autophagic compartment membranes (Jain *et al*. 2015). Aggregates of ref(2)P are considered as a marker of misfunctioning protein degradation by autophagy or the ubiquitin-proteasome system (UPS) (Bartlett *et al*. 2011; Nezis *et al*. 2008; Pircs *et al*. 2012). *Bchs* loss-of-function variants have been described to lead to a ref(2)P (human homolog: p62) accumulation and an increase in early autophagic compartments, suggesting an impaired autophagic flux (Clausen *et al*. 2010; Hebbar *et al*. 2015; Sim *et al*. 2019). To test whether *Bchs* overexpression impairs autophagy, ref(2)P immunostainings and an autophagic vesicle pH-reporter were utilized. Our data indicate strong accumulation of ref(2)P in the thoracic neuromeres of the VNC and milder also in the brain upon glial *Bchs* overexpression in the adult CNS (Fig 3A). Consistently, the overall larval CNS showed increased ref(2)P staining (Fig 2A).

In contrast, adult flies overexpressing *Bchs* panneuronally accumulated ref(2)P most prominently in the posterior region of the VNC but also with a lower signal intensity in the brain (Fig 3D). Larvae displayed ref(2)P aggregates in a subset of neurons in the VNC (Fig 2D), which we speculated to be motoneurons due to the flies’ locomotion phenotype. Driving *Bchs* overexpression in motoneurons simultaneously with a membrane-bound GFP demonstrated that a subset of motoneurons did form ref(2)P aggregates in response to *Bchs* overexpression (Fig S4). It was previously shown that *Bchs* overexpression in *even-skipped* (eve)-positive motoneurons aCC and RP2 leads to morphological abnormalities and neuronal death (Lim und Kraut 2009; Hebbar et al. 2015; Sim et al. 2019). We thus hypothesized that the ref(2)P signal might localize to eve-positive neurons, and conducted a double immunostaining of eve and ref(2)P confirming that a small subset of eve-positive neurons accumulate ref(2)P (Fig S8). However, also eve-negative neurons showed ref(2)P expression.

We further applied a GFP-mCherry-Atg8a reporter to check the ratio of non-acidic to acidic autophagic vesicles in larval brains (Nezis *et al*. 2010). Fusion of an autophagosome with an endosome or lysosome results in an acidic autophagic vesicle, termed amphisome or autolysosome, respectively. GFP fluorescence is quenched in the acidic environment leading to an mCherry-only signal that does not colocalize with a concurrent GFP signal. In larvae overexpressing *Bchs* in glial cells, a significantly increased colocalization of mCherry and GFP signals was observed in comparison to the control indicating a shift of the ratio towards non-acidic autophagic vesicles (Fig 5A,B,C,S9A). Dissimilarly, in animals with panneuronal *Bchs* overexpression no significant difference in the acidic environment of autophagic vesicles was detected (Fig 5D,E,F,S9B). Collectively, the ref(2)P accumulation and change in autophagic compartment acidity indicates that glial *Bchs* overexpression affects autophagic flux prior to or at the step of acidification of autophagic vesicles.

**Fig 5.**
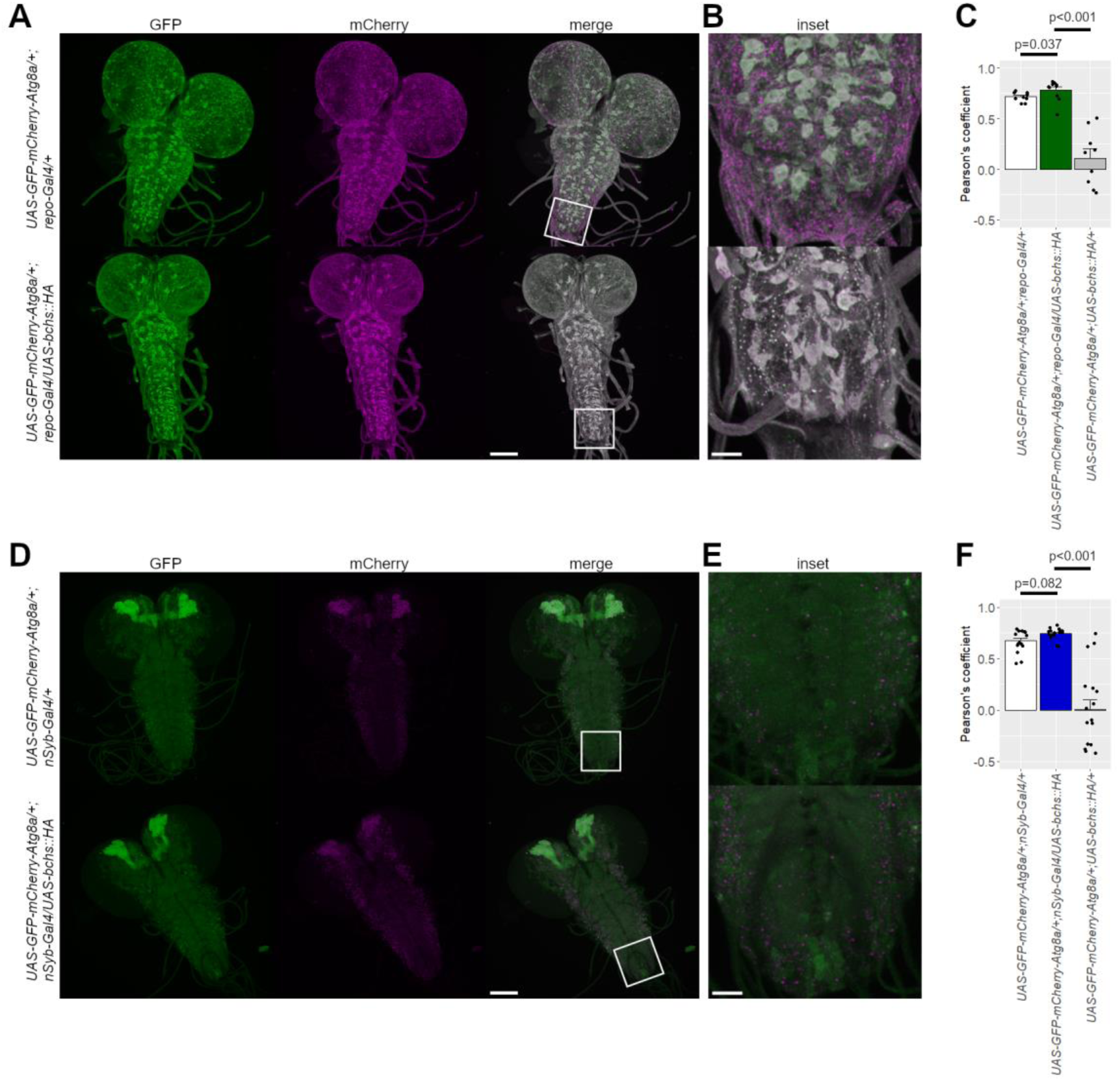
Ratio of non-acidic to acidic autophagic vesicles was altered through Bchs overexpression. A–F) The GFP-mCherry-Atg8a reporter was used to investigate autophagic flux in larval brains and was expressed in the same cell type as the *Bchs* overexpression. Autophagic flux was compared between larvae only expressing the Atg8a reporter (top, white, A−C: *UAS-GFP-mCherry-Atg8a/+; repo-Gal4/+* and D−F: *UAS-GFP-mCherry-Atg8a/+; nSyb-Gal4/+*) and larvae expressing the Atg8a reporter simultaneously with the *Bchs* overexpression (A−C: bottom, green, *UAS-GFP-mCherry-Atg8a/+; repo-Gal4/UAS-bchs::HA* and D−F: bottom, blue, *UAS-GFP-mCherry-Atg8a/+; nSyb-Gal4/UAS-bchs::HA*). As negative controls animals were used which carried both *UAS*-target genes, but not the *Gal4* (C, F: gray, *UAS-GFP-mCherry-Atg8a/+; UAS-bchs::HA/+*, Fig S9). A, D) GFP (left) and mCherry (middle) signals were imaged with a confocal microscope and images were deconvoluted. Right: GFP and mCherry images merged. Scale bar: 100 µm. Box indicates the region of the insets shown in (B) and (E). B, E) Scale bar: 20 µm. C, F) Quantification of colocalization of GFP and mCherry signals using Pearson’s coefficient. C) n ≥ 9. F) n ≥ 16. Data are shown as mean ± SEM.

### Bchs overexpression leads to altered transcriptome profiles

To understand which molecular pathways are affected by *Bchs* overexpression and how this impacts brain function, we performed RNA-seq on heads from adult flies overexpressing *Bchs* in glia or neurons. Although *Bchs* overexpression in glial cells caused a more severe phenotype with differences in brain size and autophagy when compared to neuronal overexpression (Fig 3, 5), we identified more differentially expressed genes (i.e. genes with different expression in *Bchs* overexpression flies as opposed to control animals carrying only the *Gal4* driver or only the *UAS*-target gene) in the panneuronal *nSyb-Gal4/UAS-bchs::HA* condition (2,107 genes; glial *Bchs* overexpression: 156 genes) (Tab S1). 79 genes were similarly differentially expressed upon *Bchs* overexpression in glia and neurons, which represented a highly significant overlap (*p*-value < 0.0001 from 1,000 simulations using 13,000 genes for random sampling).

Among the overlapping differentially expressed genes we identified *Iml1* and *Rab32* (Tab S1), which are known to play an important role in autophagy (Hirota & Tanaka 2009; Wang *et al*. 2012; Wu & Tu 2011). The gene ontology (GO) categories enriched for both glial and neuronal *Bchs* overexpression were related to extracellular space (Tab S1), hinting to the biological provenance of the observed phenotypes. Additionally, for glial *Bchs* overexpression the GO category septate junction (GO:0005918) was enriched. Whereas, neuronal *Bchs* overexpression led to an enrichment of GO categories related to autophagy and mitochondria.

Taken together, there was a significant number of overlapping genes that suffered dysregulation after *Bchs* overexpression in neurons or glial cells. This implies an altered molecular mechanism regardless of the inquired cell type. We identified genes involved in autophagy and lysosome formation, but also an enrichment of genes annotated to the extracellular space.

## Discussion

*WDFY3* loss-of-function variants have been linked to neurodevelopmental and neurodegenerative disorders (Dragich *et al*. 2016; Finley *et al*. 2003; Le Duc *et al*. 2019; Orosco *et al*. 2014; Schaaf *et al*. 2022). Further, *WDFY3* variants have been associated with an increase or decrease in brain size (Finley *et al*. 2003; Kadir *et al*. 2016; Kriston-Vizi *et al*. 2011; Le Duc *et al*. 2019; Orosco *et al*. 2014). We hypothesized that variants underlying potential loss– or gain-of-function have opposing effects on brain size. Here, we used a *Drosophila Bchs* overexpression model as a proxy for gain-of-function effects in development. Ubiquitous *Bchs* overexpression was lethal (Fig S1), suggesting that *Bchs* expression levels are highly relevant also in non-neural tissue. We then overexpressed *Bchs* in different cell types of the nervous system to understand how dysregulation in different cells impacts development.

### Bchs overexpression impaired nervous system, wing, and thorax development

We tested whether dysregulation of *WDFY3*/*Bchs* in glial cells may contribute to phenotypic abnormalities observed in probands with *WDFY3* variants. We detected a prolonged developmental time for flies overexpressing *Bchs* in glial cells, but not for panneuronal overexpression, indicating a role of glial *Bchs* in developmental processes (Fig 1A).

On the other side, neuronal *Bchs* overexpression caused higher rates of wing and thorax defects. Driving *Bchs* overexpression only in CCAP neurons was sufficient to cause these morphological phenotypes, implicating that CCAP neurons are responsible for wing and thorax abnormalities. CCAP neurons are well known to have a major role in post-ecdysis development (Luan *et al*. 2006; Park *et al*. 2003). Suppressing their activity disrupts tonic abdominal contractions and air swallowing, a motor program necessary to pump hemolymph into wings to unfold them (Peabody *et al*. 2009). Further, CCAP neurons secrete the hormone bursicon, required for wing expansion and tanning, into the hemolymph (Dewey *et al*. 2004; Loveall & Deitcher 2010; Luo *et al*. 2005). Interestingly, similar wing and thorax abnormalities were noticed upon misexpression of *TBPH* (human ortholog: *TDP-43*), an RNA-binding protein, or knockdown of *Gclc*, an enzyme involved in glutathione synthesis, in CCAP neurons (Mercer *et al*. 2016; Vanden Broeck *et al*. 2013). Dysregulations of those genes were hypothesized to induce premature degeneration of CCAP neurons causing wing and thorax defects. Therefore, we suspect that also *Bchs* overexpression might have contributed to a degeneration of CCAP neurons. Importantly, WDFY3 is involved in the removal of mutant TDP-43 (Han *et al*. 2015), suggesting that *Bchs* overexpression might lead to a misregulation of TBPH in CCAP neurons. However, in our transcriptomic analyses, we did not identify *TBPH* or *Gclc* to be differentially expressed in flies overexpressing *Bchs* in neurons. On the other side, *Pburs*, a subunit of the hormone bursicon was upregulated (Tab S1, 10-fold higher expression, *adj-p*= 0.0008) (Luo *et al*. 2005). Null mutants of *Pburs* were similarly described to have a wing expansion deficit (Lahr *et al*. 2012). The observed weak penetrance of wing and thorax abnormalities in glial *Bchs* overexpression flies may indicate that in a small number of flies glial *Bchs* overexpression provoked a misfunctioning of CCAP neurons (Fig 1D,E). Similarly, the weak penetrance of abnormalities in motoneuronal *Bchs* overexpression could be explained by its expression in CCAP motoneurons but not in CCAP interneurons (Fig S3). Our data suggest that both, neuronal and glial *Bchs* dysregulation contribute to developmental defects. This is further supported by the impaired crawling behavior in glial and neuronal *Bchs* overexpression larvae (Fig 2C,F), which is in accordance with delayed motor development observed in affected patients (Le Duc *et al*. 2019).

### Bchs affects CNS morphology

*WDFY3* loss-of-function variants lead to an increased brain size in mice and humans (Le Duc *et al*. 2019; Orosco *et al*. 2014), while *Bchs* loss-of-function variants were described to reduce the brain size (Finley *et al*. 2003; Kriston-Vizi *et al*. 2011). Although dysregulated *WDFY3/Bchs* levels might have different downstream effects in the respective model organisms, we were still prompted to check whether the overexpression in the different nervous system cells impact the overall brain size.

In our study, only glial, but not neuronal, *Bchs* overexpression caused alterations in CNS size, presented in an elongated VNC and a decreased brain size to fly length ratio (Fig 2,3). In contrast, previous studies demonstrated that neuronal *Bchs* overexpression increases larval brain size (Kriston-Vizi *et al*. 2011). This discrepancy of neuronal overexpression phenotypes might be due to investigation of different developmental stages or because of using different neuronal drivers. Importantly, glial *Bchs* can influence the CNS size, demonstrating that glia need to be considered when examining the mechanism underlying altered CNS size. Furthermore, different cell types also of distinct stages of differentiation might contribute to the altered CNS size of *WDFY3*/*Bchs* mutants. The contribution of each cell type might depend on the observed developmental phase. In mice, Orosco *et al*. showed that hypomorphic variants of *Wdfy3* increase symmetric proliferation of radial glia, neural stem cells, which give rise to neurons and glia (Orosco *et al*. 2014). Interestingly, glial *Bchs* overexpression resulted in a gain of glial cell number (Fig 4,S7). This gain could have been caused similarly by increased proliferation of *Drosophila* neural stem cells (neuroblasts), intermediate progenitor cells or glial cells. It is well known that glial cells can regulate neuroblast proliferation (Contreras *et al*. 2021; Kanai *et al*. 2018; Nguyen & Cheng 2022; Yang *et al*. 2021). However, reduced glial cell death could also have provoked the cell number increase. Further studies are needed, to determine whether the increased glial cell number is caused by alteration of proliferation or cell death.

Several genes have been previously associated with elongated VNC as seen in glial *Bchs* overexpression larvae (Fig 2), e.g. glial overexpression of the genes *mmp2* or *kuz,* encoding metalloproteases (Dai *et al*. 2018; Kato *et al*. 2011; Losada-Perez *et al*. 2016; Meyer *et al*. 2014; Pandey *et al*. 2011; Skeath *et al*. 2017; Winkler *et al*. 2021). Both proteases likely regulate BBB integrity (Kanda *et al*. 2019; Petri *et al*. 2019). Subperineural glia form an essential part of the BBB by producing septate junctions. Importantly, *Bchs* overexpression in subperineural glia was sufficient to promote VNC elongation. Additionally, in our transcriptomic analyses we found a dysregulation of genes associated with septate junctions (*Tsf2*, *kune*, *cold*, *gli*, *hoka*, *udt*, Tab S1) in flies overexpressing *Bchs* in glial cells (Hijazi *et al*. 2011; Izumi *et al*. 2021; Kanda *et al*. 2019; Nelson *et al*. 2010; Schulte *et al*. 2003; Tiklová *et al*. 2010). This suggests that glial *Bchs* overexpression impairs proper septate junction formation and BBB integrity, which could contribute to the altered brain size observed in different animal models and probands.

### Bchs overexpression impairs autophagic flux

Further, transcriptomics analyses identified 2,107 and 156 genes to be differentially expressed in flies overexpressing *Bchs* in neurons and glia, respectively (Tab S1). For neuronal and glial *Bchs* overexpression, 50 and 3 of the differentially expressed genes, respectively, are annotated to the GO category autophagy (GO:0006914) or its child terms. Those genes play important roles in autophagy, which is the main known function of *WDFY3*/*Bchs*.

Bchs is an adaptor between ref(2)P and the autophagosomal membrane, therefore, a gain of *Bchs* could be assumed to increase autophagic flux (Clausen *et al*. 2010; Filimonenko *et al*. 2010; Sim *et al*. 2019; Simonsen *et al*. 2004). However, our data showing ref(2)P aggregation and a shifted ratio towards non-acidic autophagic vesicles when *Bchs* was overexpressed in glia suggest that the overexpression disrupts the autophagy pathway (Fig 2,3,5). Additionally, neuronal and glial *Bchs* overexpression decreased mRNA levels of the autophagy-associated genes *Iml1* and *Rab32* (*lightoid*) (Tab S1), further supporting a perturbed autophagic flux (Wang *et al*. 2012; Wu & Tu 2011). Nevertheless, we cannot rule out that formation of non-acidic autophagic vesicles was increased but the downstream autophagy pathway was limited by the fusion step, leading to an excessive build-up of non-acidic vesicles.

In both, larval and adult CNS, glial *Bchs* overexpression compared to neuronal *Bchs* overexpression resulted in a ref(2)P aggregation pattern that was more widely spread in the CNS (Fig 2,3). However, from ref(2)P accumulation it is not possible to conclude on the impact on neuronal function. Yet, recent research has shown that glial autophagy impacts neuronal health, e.g. in neurodegenerative diseases like Parkinson’s disease and Alzheimer’s disease (Bankston *et al*. 2019; Cho *et al*. 2014; Choi *et al*. 2020; Damulewicz *et al*. 2022; Kreher *et al*. 2021; Szabó *et al*. 2023; Tu *et al*. 2021).

In this study, we demonstrated that glial, as well as neuronal *Bchs* overexpression can lead to developmental abnormalities. While neurons have been implicated in *WDFY3*-associated pathologies, our data indicate that glial *Bchs* dysregulation also contributes to phenotypic defects. Since at least in respect to brain size *WDFY3*/*Bchs* was shown to yield different downstream effects in different model organisms, further investigations should be carried out to decipher the role of glial *WDFY3* in *WDFY3*-related neurodevelopmental disorders, and whether a modulation of glial function could rescue the phenotype.

## Materials and Methods

### Fly husbandry

Flies were maintained on standard cornmeal food at 25 °C and a 12:12 light-dark cycle. The following *Drosophila melanogaster* strains were used:

*_w_1118*

*;; repo-GAL4/TM6B Tb*, (RRID:BDSC_7415)

*;; nSyb-GAL4/Sb*, (gift from J. Simpson)

*; UAS-bchs::HA*, (RRID:BDSC_51636)

*;; act5C-Gal4/Tb,Sb*, (RRID:BDSC_3954)

*; ok6-GAL4 w^+^*, (Marqués *et al*. 2002)

*; UASp-GFP-mCherry-Atg8a*, (RRID:BDSC_37749)

*; UAS-mCD8::GFP, UAS-mCD8::GFP*, (RRID:BDSC_5137)

*;; UAS-mCD8::GFP*, (RRID:BDSC_5130)

*; Burs-GAL4*, (RRID:BDSC_40972)

*;; CCAP-GAL4*, (RRID:BDSC_25686)

*; rL82-GAL4*, (Sepp & Auld 1999)

*; nrv2-GAL4*, (RRID:BDSC_6800)

*;; c527-Gal4*, (RRID:BDSC_90391)

### Mendelian ratio and adult developmental time

15 virgin females (*UAS-bchs::HA/Sb^1^*) and 5 males (cell type-specific reporter-*Gal4/Sb^1^*), all carrying the balancer chromosome (marker: *Sb^1^*), were crossed and switched to a new vial every day. Numbers of flies carrying the balancer chromosome (no *Bchs* overexpression) and flies not carrying the balancer (*Bchs* overexpression) in the F1 generation of five vials were counted for each genotype. Newly hatched flies were counted every 24 h. For each vial, counting was started on the day the first adult fly hatched and continued for ten days.

### PEDtracker – embryonal developmental time, larval and pupal size

The development of panneuronal *Bchs* overexpression flies (*nSyb-Gal4/UAS-bchs::HA*) as well as the controls *nSyb-Gal4/+* and *UAS-bchs::HA/+* was monitored using the PEDtracker system (Schumann & Triphan 2020). According to the previously published protocol, the development of the specimen was observed from egg hatching to pupation. Larval hatching timepoint, larval size over development as well as pupal size were then manually measured using ImageJ (Schneider *et al*. 2012). Statistical analysis was performed using Kruskal-Wallis test or one-way ANOVA depending on the distribution of the data.

### Assessing the condition of wings and thoraces

20 virgin females and 10 males were crossed and switched to a new vial every second day. For each genotype, the F1 generation in three vials was analysed. For each vial, collecting flies was started on the day the first adult fly hatched and continued for five days. The condition of wings and thoraces was evaluated 24 h after collecting the flies (24−48 h old flies). Wings were categorized into folded, partially folded, and expanded. Thoraces were categorized into dimpled and not dimpled.

### Larval locomotion

Petri dishes with a diameter of 9 cm filled with 1 % agarose were prepared. Five third instar larvae were placed in the middle of a petri dish and recorded (camera: Logitech C920 HD Pro) for 1 min. The first 5 sec of a recording were dismissed and larval behavior of the following 30 sec investigated. Recordings were analysed by using the ImageJ freehand line tool to measure the length of the crawled route of a larva. For each genotype, 30 larvae were examined if not stated otherwise.

### Immunohistochemistry

Larvae: Wandering third instar larvae were dissected in ice-cold HL-3 solution (Stewart *et al*. 1994), fixed with PFA (4 % in PBS) and collected in 1× PBS. Blocking, primary antibody incubation and secondary antibody incubation were consecutively performed in PBT (1× PBS + 0.05 % Triton X-100, Sigma-Aldrich) containing 5 % normal goat serum (NGS, Jackson ImmunoResearch) at 4°C overnight. Antibody incubation steps were followed by washing two times shortly and three times 15 min (1× PBS + 0.05 % Triton X-100). Samples were stored in Vectashield (Vector Laboratories) at 4°C overnight before mounting.

Adult flies: The CNS of adult female flies (21−33 h old) were dissected in ice-cold Ringer solution, fixed in 4 % PFA for 30 min at room temperature and collected in 1× PBS. Blocking was done at room temperature for 24 h in PBT (1× PBS + 1 % Triton X-100) containing 5 % NGS. Primary antibody incubation was carried out at 4°C for 24 h and secondary antibody incubation at 4°C for 24−48 h. Antibody incubation steps were followed by moving samples to room temperature for 1 h and washing two times shortly and three times 15 min with PBT. Samples were stored in Vectashield at 4°C overnight before mounting.

The following antibodies were used at following dilutions: rabbit-anti-Ref(2)P (1:500, Abcam, ab178440), mouse-anti-repo (1:250, DSHB, 8D12 concentrate), mouse-anti-even skipped (1:50, DSHB, 2B8), goat-anti-mouse conjugated with Alexa Fluor-488 or –405 (1:250, Invitrogen, A-11001, RRID: AB_2534069 and A-31553, RRID: AB_221604), goat-anti-horseradish peroxidase (HRP) conjugated with Alexa Fluor-488 or –647 (1:250, Jackson ImmunoResearch, 123-545-021, RRID: AB_2338965 and 123-605-021, RRID: AB_2338967), Cy3-or Cy5-conjugated goat-anti-rabbit (1:250, Jackson ImmunoResearch, 111-165-144, RRID:AB_2338006 and 111-175-144, RRID: AB_2338013).

### Central nervous system size measurement

Larval VNC was measured and normalized to the larval length which was determined before dissection. Larvae were placed into a petri dish filled with HL-3 on ice and imaged (camera: Leica DFC365 FX, microscope: Leica MZ10 F). Brains were stained against HRP, imaged and a maximum projection of the z-stack was performed with ImageJ. The length of the larvae and the VNC were measured with the ImageJ Straight Line tool.

The size of the VNC and the brain of adult flies was determined and normalized to the fly length. The adult fly was anesthetized by putting it in a vial on ice. The fly was transferred to a petri dish and imaged (camera: Leica DFC365 FX, microscope: Leica MZ10 F). The central nervous system was stained against HRP, imaged, and a maximum projection was performed. Lengths of the fly and the VNC were measured with the ImageJ Straight Line tool. The area of the brain was measured with the ImageJ Freehand selection tool.

### Number of glial cells

Larvae were stained with anti-repo antibody and imaged. Repo-positive nuclei were counted using the ImageJ plugin cell counter. For analysis of glial cell number in the peripheral nervous system, glial nuclei number was determined at the entry point of the peripheral nerve bundle into the body wall muscles at the region, where the bundle divides into the TN, ISN, and SN nerve branches. Peripheral nerves innervating the hemisegments A4R and A4L were analysed.

### Colocalization of GFP-mCherry-Atg8 fluorophore signals

Larval brains were dissected and fixed as described before, stored in Vectashield for at least 24 h and imaged. Images were deconvoluted using Huygens Essentials software (strategy: standard). Analyses of images were performed with the ImageJ plugin coloc 2 (Pearson correlation, above threshold) and using the freehand selection tool to set a ROI surrounding the brain.

### Imaging

Image acquisition was performed with a Leica SP8 confocal microscope unless specified otherwise.

### RNA extraction and sequencing

RNA was extracted from five female and five male fly heads for one sample. Flies (22−31 h after eclosion) were anesthetized with carbon dioxide and heads were cut using a scalpel. Heads were immediately transferred to an ice-cold 2 ml Eppendorf tube containing Trizol or RLT buffer. RNA extraction followed immediately using the RNeasy Micro Kit (Qiagen, Cat. No. 74004) according to protocol or exchanging the first step of homogenization in RLT buffer with the following Trizol protocol (Invitrogen, Cat. No. 15596026). Heads were homogenized in 500 µl Trizol, centrifuged for 5 min at 10,000 rpm at 4°C, supernatant was tranferred to a new tube, 200 µl chloroform (Carl Roth, No. 6340.1) was added and the tube was shaken for 15 sec. An incubation step at room temperature for 3 min was followed by centrifugation for 5 min at 10,000 rpm at 4°C. The upper aqueous phase was transferred in a new tube and it was continued with the RNeasy Micro Kit protocol. Homogenization was performed on ice using the Ultra-Turrax (IKA T10 basic) five times for 10 sec. For each genotype five samples were sequenced.

RNA-seq libraries were prepared using TruSeq RNA Library Prep Kit v2 (Illumina, San Diego, CA) and sequenced on an Illumina NovaSeq platform with 151 bp paired-end reads with an average of ∼135 million reads per library.

### Differential Gene Expression (DEG) analysis

RNA-Seq reads were mapped to the Drosophila genome assembly BDGP6.32 (GCA_000001215.4) with STAR (version 2.6.1d) (Dobin *et al*. 2013). Reads were processed as previously described (Körner *et al*. 2022). We computed the transcript levels with htseq-count (version 0.6.0) (Anders *et al*. 2015). Genes with a sum of less than 10 reads in all samples together were excluded from further analysis. Differential expression of genes was determined with the R package DESeq2 (version 1.30.1) (Love *et al*. 2014), which uses the Benjamini-Hochberg method to correct for multiple testing (Benjamini *et al*. 2001). Genes were considered to be significantly differentially expressed if *p*-adj < 0.05. To check clustering of RNA-sequencing samples of subjects and controls, a principal component analysis (PCA) was performed with the R package pcaExplorer (version 2.6.0) (Marini und Binder 2019). RNA count data were variance stabilized transformed and the 500 most variable genes (top n genes) were selected for computing the principal components. We tested 5 samples of adult heads from glial/neuronal *Bchs* overexpression (*repo-Gal4/UAS-bchs::HA* and *nSyb-Gal4/UAS-bchs::HA*, respectively) against the pooled samples of both controls, 5 samples of the *Gal4*-driver control (*repo-Gal4/+* and *nSyb-Gal4/+*, respectively) and 5 samples of the *UAS* control (*UAS-bchs::HA/+*).

To identify which pathways are enriched with differentially expressed genes we used the GOfuncR package with a hypergeometric test and the Drosophila GO annotations org.Dm.eg.db v.3.17 (Carlson 2019; Grote 2021).

### Statistics

Data are shown as mean ± SEM. Statistical analyses were performed with SigmaPlot 12.5 (Systat software) using two-tailed Student’s t-tests or Mann-Whitney Rank Sum test for non-normally distributed data, if not stated otherwise.

## Acknowledgments

We thank Anne Butthof for advices on RNA extraction protocol, Mathias Böhme for discussion about glial cells and Torsten Schöneberg for useful discussions which led to new insights in respect to the pathomechanism.

## Author contributions

MK performed fly experiments, contributed to the experimental design and wrote the first draft of the manuscript. AV and LB performed gene expression and GO enrichment analyses. IS and JK performed PEDtracker assays and contributed to writing of the manuscript. KK, KD and JH performed RNA and DNA sequencing. AG, AT and RAJ contributed to result interpretation and writing of the manuscript. NS, AM and TL contributed to design of the study, data interpretation, funding acquisition, and writing of the manuscript. DLD designed and supervised the study, acquired funding, and contributed to writing of the manuscript.

## Availability of data and materials

RNA-seq data has been submitted to the Gene Expression Omnibus (http://www.ncbi.nlm.nih.gov/geo/) under accession number GSE244775.

## References

1. Aber ER, Griffey CJ, Davies T, Li AM, Yang YJ, Croce KR, Goldman JE, Grutzendler J, Canman JC & Yamamoto A (2022). Oligodendroglial macroautophagy is essential for myelin sheath turnover to prevent neurodegeneration and death, Cell Rep 41, 111480; DOI: 10.1016/j.celrep.2022.111480.

2. Anders S, Pyl PT & Huber W (2015). HTSeq—a Python framework to work with high-throughput sequencing data, Bioinformatics 31, 166–169; DOI: 10.1093/bioinformatics/btu638.

3. Bankston AN, Forston MD, Howard RM, Andres KR, Smith AE, Ohri SS, Bates ML, Bunge MB & Whittemore SR (2019). Autophagy is essential for oligodendrocyte differentiation, survival, and proper myelination, Glia 67, 1745–1759; DOI: 10.1002/glia.23646.

4. Bartlett BJ, Isakson P, Lewerenz J, Sanchez H, Kotzebue RW, Cumming RC, Harris GL, Nezis IP, Schubert DR & Simonsen A, et al. (2011). p62, Ref(2)P and ubiquitinated proteins are conserved markers of neuronal aging, aggregate formation and progressive autophagic defects, Autophagy 7, 572–583; DOI: 10.4161/auto.7.6.14943.

5. Baumgartner S, Littleton JT, Broadie K, Bhat MA, Harbecke R, Lengyel JA, Chiquet-Ehrismann R, Prokop A & Bellen HJ (1996). A Drosophila neurexin is required for septate junction and blood-nerve barrier formation and function, Cell 87, 1059–1068; DOI: 10.1016/s0092-8674(00)81800-0.

6. Benjamini Y, Drai D, Elmer G, Kafkafi N & Golani I (2001). Controlling the false discovery rate in behavior genetics research, Behav Brain Res 125, 279–284; DOI: 10.1016/s0166-4328(01)00297-2.

7. Bittern J, Pogodalla N, Ohm H, Brüser L, Kottmeier R, Schirmeier S & Klämbt C (2021). Neuron-glia interaction in the Drosophila nervous system, Dev Neurobiol 81, 438–452; DOI: 10.1002/dneu.22737.

8. Carlson M (2019). org.Dm.eg.db: Genome wide annotation for Fly. R package version 3.8.2.

9. Cho M-H, Cho K, Kang H-J, Jeon E-Y, Kim H-S, Kwon H-J, Kim H-M, Kim D-H & Yoon S-Y (2014). Autophagy in microglia degrades extracellular β-amyloid fibrils and regulates the NLRP3 inflammasome, Autophagy 10, 1761–1775; DOI: 10.4161/auto.29647.

10. Choi I, Zhang Y, Seegobin SP, Pruvost M, Wang Q, Purtell K, Zhang B & Yue Z (2020). Microglia clear neuron-released α-synuclein via selective autophagy and prevent neurodegeneration, Nat Commun 11, 1386; DOI: 10.1038/s41467-020-15119-w.

11. Clausen TH, Lamark T, Isakson P, Finley K, Larsen KB, Brech A, Øvervatn A, Stenmark H, Bjørkøy G & Simonsen A, et al. (2010). p62/SQSTM1 and ALFY interact to facilitate the formation of p62 bodies/ALIS and their degradation by autophagy, Autophagy 6, 330–344; DOI: 10.4161/auto.6.3.11226.

12. Contreras EG, Glavic Á, Brand AH & Sierralta JA (2021). The Serine Protease Homolog, Scarface, Is Sensitive to Nutrient Availability and Modulates the Development of the Drosophila Blood-Brain Barrier, J. Neurosci. 41, 6430–6448; DOI: 10.1523/JNEUROSCI.0452-20.2021.

13. Dai J, Estrada B, Jacobs S, Sánchez-Sánchez BJ, Tang J, Ma M, Magadán-Corpas P, Pastor-Pareja JC & Martín-Bermudo MD (2018). Dissection of Nidogen function in Drosophila reveals tissue-specific mechanisms of basement membrane assembly, PLoS Genet 14, e1007483; DOI: 10.1371/journal.pgen.1007483.

14. Damulewicz M, Szypulski K & Pyza E (2022). Glia-Neurons Cross-Talk Regulated Through Autophagy, Front Physiol 13, 886273; DOI: 10.3389/fphys.2022.886273.

15. Dewey EM, McNabb SL, Ewer J, Kuo GR, Takanishi CL, Truman JW & Honegger H-W (2004). Identification of the gene encoding bursicon, an insect neuropeptide responsible for cuticle sclerotization and wing spreading, Curr Biol 14, 1208–1213; DOI: 10.1016/j.cub.2004.06.051.

16. DiAntonio A, Burgess RW, Chin AC, Deitcher DL, Scheller RH & Schwarz TL (1993). Identification and characterization of Drosophila genes for synaptic vesicle proteins, J. Neurosci. 13, 4924–4935; DOI: 10.1523/JNEUROSCI.13-11-04924.1993.

17. Dobin A, Davis CA, Schlesinger F, Drenkow J, Zaleski C, Jha S, Batut P, Chaisson M & Gingeras TR (2013). STAR: ultrafast universal RNA-seq aligner, Bioinformatics 29, 15– 21; DOI: 10.1093/bioinformatics/bts635.

18. Dragich JM, Kuwajima T, Hirose-Ikeda M, Yoon MS, Eenjes E, Bosco JR, Fox LM, Lystad AH, Oo TF & Yarygina O, et al. (2016). Autophagy linked FYVE (Alfy/WDFY3) is required for establishing neuronal connectivity in the mammalian brain, eLife 5; DOI: 10.7554/eLife.14810.

19. Filimonenko M, Isakson P, Finley KD, Anderson M, Jeong H, Melia TJ, Bartlett BJ, Myers KM, Birkeland HCG & Lamark T, et al. (2010). The selective macroautophagic degradation of aggregated proteins requires the PI3P-binding protein Alfy, Mol Cell 38, 265–279; DOI: 10.1016/j.molcel.2010.04.007.

20. Finley KD, Edeen PT, Cumming RC, Mardahl-Dumesnil MD, Taylor BJ, Rodriguez MH, Hwang CE, Benedetti M & McKeown M (2003). blue cheese Mutations Define a Novel, Conserved Gene Involved in Progressive Neural Degeneration, J. Neurosci. 23, 1254– 1264; DOI: 10.1523/JNEUROSCI.23-04-01254.2003.

21. Fox LM, Kim K, Johnson CW, Chen S, Croce KR, Victor MB, Eenjes E, Bosco JR, Randolph LK & Dragatsis I, et al. (2020). Huntington’s Disease Pathogenesis Is Modified In Vivo by Alfy/Wdfy3 and Selective Macroautophagy, Neuron 105, 813–821.e6; DOI: 10.1016/j.neuron.2019.12.003.

22. Grote S (2021). GOfuncR: Gene ontology enrichment using FUNC. R package version 1.14.0.

23. Han H, Wei W, Duan W, Guo Y, Li Y, Wang J, Bi Y & Li C (2015). Autophagy-linked FYVE protein (Alfy) promotes autophagic removal of misfolded proteins involved in amyotrophic lateral sclerosis (ALS), In Vitro Cell Dev Biol Anim 51, 249–263; DOI: 10.1007/s11626-014-9832-4.

24. Hebbar S, Sahoo I, Matysik A, Argudo Garcia I, Osborne KA, Papan C, Torta F, Narayanaswamy P, Fun XH & Wenk MR, et al. (2015). Ceramides And Stress Signalling Intersect With Autophagic Defects In Neurodegenerative Drosophila blue cheese (bchs) Mutants, Sci Rep 5, 15926; DOI: 10.1038/srep15926.

25. Hijazi A, Haenlin M, Waltzer L & Roch F (2011). The Ly6 protein coiled is required for septate junction and blood brain barrier organisation in Drosophila, PLoS ONE 6, e17763; DOI: 10.1371/journal.pone.0017763.

26. Hirota Y & Tanaka Y (2009). A small GTPase, human Rab32, is required for the formation of autophagic vacuoles under basal conditions, Cell Mol Life Sci 66, 2913–2932; DOI: 10.1007/s00018-009-0080-9.

27. Izumi Y, Furuse K & Furuse M (2021). The novel membrane protein Hoka regulates septate junction organization and stem cell homeostasis in the Drosophila gut, J Cell Sci 134; DOI: 10.1242/jcs.257022.

28. Jain A, Rusten TE, Katheder N, Elvenes J, Bruun J-A, Sjøttem E, Lamark T & Johansen T (2015). p62/Sequestosome-1, Autophagy-related Gene 8, and Autophagy in Drosophila Are Regulated by Nuclear Factor Erythroid 2-related Factor 2 (NRF2), Independent of Transcription Factor TFEB, J Biol Chem 290, 14945–14962; DOI: 10.1074/jbc.M115.656116.

29. Kadir R, Harel T, Markus B, Perez Y, Bakhrat A, Cohen I, Volodarsky M, Feintsein-Linial M, Chervinski E & Zlotogora J, et al. (2016). ALFY-Controlled DVL3 Autophagy Regulates Wnt Signaling, Determining Human Brain Size, PLoS Genet 12, e1005919; DOI: 10.1371/journal.pgen.1005919.

30. Kanai MI, Kim M-J, Akiyama T, Takemura M, Wharton K, O’Connor MB & Nakato H (2018). Regulation of neuroblast proliferation by surface glia in the Drosophila larval brain, Sci Rep 8, 3730; DOI: 10.1038/s41598-018-22028-y.

31. Kanda H, Shimamura R, Koizumi-Kitajima M & Okano H (2019). Degradation of Extracellular Matrix by Matrix Metalloproteinase 2 Is Essential for the Establishment of the Blood-Brain Barrier in Drosophila, iScience 16, 218–229; DOI: 10.1016/j.isci.2019.05.027.

32. Karlsson M, Zhang C, Méar L, Zhong W, Digre A, Katona B, Sjöstedt E, Butler L, Odeberg J & Dusart P, et al. (2021). A single-cell type transcriptomics map of human tissues, Sci Adv 7; DOI: 10.1126/sciadv.abh2169.

33. Kato K, Forero MG, Fenton JC & Hidalgo A (2011). The glial regenerative response to central nervous system injury is enabled by pros-notch and pros-NFκB feedback, PLoS Biol 9, e1001133; DOI: 10.1371/journal.pbio.1001133.

34. Khodosh R, Augsburger A, Schwarz TL & Garrity PA (2006). Bchs, a BEACH domain protein, antagonizes Rab11 in synapse morphogenesis and other developmental events, Development 133, 4655–4665; DOI: 10.1242/dev.02650.

35. Kim YS, Choi J & Yoon B-E (2020). Neuron-Glia Interactions in Neurodevelopmental Disorders, Cells 9; DOI: 10.3390/cells9102176.

36. Körner MB, Velluva A, Bundalian L, Radtke M, Lin C-C, Zacher P, Bartolomaeus T, Kirstein AS, Mrestani A & Scholz N, et al. (2022). Altered gene expression profiles impair the nervous system development in individuals with 15q13.3 microdeletion, Sci Rep 12, 13507; DOI: 10.1038/s41598-022-17604-2.

37. Kraut R, Menon K & Zinn K (2001). A gain-of-function screen for genes controlling motor axon guidance and synaptogenesis in Drosophila, Curr Biol 11, 417–430; DOI: 10.1016/S0960-9822(01)00124-5.

38. Kreher C, Favret J, Maulik M & Shin D (2021). Lysosomal Functions in Glia Associated with Neurodegeneration, Biomolecules 11; DOI: 10.3390/biom11030400.

39. Kriston-Vizi J, Thong NW, Poh CL, Yee KC, Ling JSP, Kraut R & Wasser M (2011). Gebiss: an ImageJ plugin for the specification of ground truth and the performance evaluation of 3D segmentation algorithms, BMC Bioinformatics 12, 232; DOI: 10.1186/1471-2105-12-232.

40. Lago-Baldaia I, Fernandes VM & Ackerman SD (2020). More Than Mortar: Glia as Architects of Nervous System Development and Disease, Front Cell Dev Biol 8, 611269; DOI: 10.3389/fcell.2020.611269.

41. Lahr EC, Dean D & Ewer J (2012). Genetic analysis of ecdysis behavior in Drosophila reveals partially overlapping functions of two unrelated neuropeptides, J. Neurosci. 32, 6819–6829; DOI: 10.1523/JNEUROSCI.5301-11.2012.

42. Le Duc D, Giulivi C, Hiatt SM, Napoli E, Panoutsopoulos A, Harlan De Crescenzo A, Kotzaeridou U, Syrbe S, Anagnostou E & Azage M, et al. (2019). Pathogenic WDFY3 variants cause neurodevelopmental disorders and opposing effects on brain size, Brain 142, 2617–2630; DOI: 10.1093/brain/awz198.

43. Lim A & Kraut R (2009). The Drosophila BEACH family protein, blue cheese, links lysosomal axon transport with motor neuron degeneration, J. Neurosci. 29, 951–963; DOI: 10.1523/JNEUROSCI.2582-08.2009.

44. Losada-Perez M, Harrison N & Hidalgo A (2016). Molecular mechanism of central nervous system repair by the Drosophila NG2 homologue kon-tiki, J Cell Biol 214, 587–601; DOI: 10.1083/jcb.201603054.

45. Love MI, Huber W & Anders S (2014). Moderated estimation of fold change and dispersion for RNA-seq data with DESeq2, Genome Biol 15, 550; DOI: 10.1186/s13059-014-0550-8.

46. Loveall BJ & Deitcher DL (2010). The essential role of bursicon during Drosophila development, BMC Dev Biol 10, 92; DOI: 10.1186/1471-213X-10-92.

47. Luan H, Lemon WC, Peabody NC, Pohl JB, Zelensky PK, Wang D, Nitabach MN, Holmes TC & White BH (2006). Functional dissection of a neuronal network required for cuticle tanning and wing expansion in Drosophila, J. Neurosci. 26, 573–584; DOI: 10.1523/JNEUROSCI.3916-05.2006.

48. Luo C-W, Dewey EM, Sudo S, Ewer J, Hsu SY, Honegger H-W & Hsueh AJW (2005). Bursicon, the insect cuticle-hardening hormone, is a heterodimeric cystine knot protein that activates G protein-coupled receptor LGR2, Proc Natl Acad Sci U S A 102, 2820–2825; DOI: 10.1073/pnas.0409916102.

49. Marqués G, Bao H, Haerry TE, Shimell MJ, Duchek P, Zhang B & O’Connor MB (2002). The Drosophila BMP type II receptor Wishful Thinking regulates neuromuscular synapse morphology and function, Neuron 33, 529–543; DOI: 10.1016/s0896-6273(02)00595-0.

50. Mercer SW, La Fontaine S, Warr CG & Burke R (2016). Reduced glutathione biosynthesis in Drosophila melanogaster causes neuronal defects linked to copper deficiency, J Neurochem 137, 360–370; DOI: 10.1111/jnc.13567.

51. Meyer S, Schmidt I & Klämbt C (2014). Glia ECM interactions are required to shape the Drosophila nervous system, Mech Dev 133, 105–116; DOI: 10.1016/j.mod.2014.05.003.

52. Nelson KS, Furuse M & Beitel GJ (2010). The Drosophila Claudin Kune-kune is required for septate junction organization and tracheal tube size control, Genetics 185, 831–839; DOI: 10.1534/genetics.110.114959.

53. Nezis IP, Shravage BV, Sagona AP, Lamark T, Bjørkøy G, Johansen T, Rusten TE, Brech A, Baehrecke EH & Stenmark H (2010). Autophagic degradation of dBruce controls DNA fragmentation in nurse cells during late Drosophila melanogaster oogenesis, J Cell Biol 190, 523–531; DOI: 10.1083/jcb.201002035.

54. Nezis IP, Simonsen A, Sagona AP, Finley K, Gaumer S, Contamine D, Rusten TE, Stenmark H & Brech A (2008). Ref(2)P, the Drosophila melanogaster homologue of mammalian p62, is required for the formation of protein aggregates in adult brain, J Cell Biol 180, 1065–1071; DOI: 10.1083/jcb.200711108.

55. Nguyen P-K & Cheng LY (2022). Non-autonomous regulation of neurogenesis by extrinsic cues: a Drosophila perspective, Oxf Open Neurosci 1; DOI: 10.1093/oons/kvac004.

56. Orosco LA, Ross AP, Cates SL, Scott SE, Wu D, Sohn J, Pleasure D, Pleasure SJ, Adamopoulos IE & Zarbalis KS (2014). Loss of Wdfy3 in mice alters cerebral cortical neurogenesis reflecting aspects of the autism pathology, Nat Commun 5, 4692; DOI: 10.1038/ncomms5692.

57. Pandey R, Blanco J & Udolph G (2011). The glucuronyltransferase GlcAT-P is required for stretch growth of peripheral nerves in Drosophila, PLoS ONE 6, e28106; DOI: 10.1371/journal.pone.0028106.

58. Park JH, Schroeder AJ, Helfrich-Förster C, Jackson FR & Ewer J (2003). Targeted ablation of CCAP neuropeptide-containing neurons of Drosophila causes specific defects in execution and circadian timing of ecdysis behavior, Development 130, 2645–2656; DOI: 10.1242/dev.00503.

59. Peabody NC, Pohl JB, Diao F, Vreede AP, Sandstrom DJ, Wang H, Zelensky PK & White BH (2009). Characterization of the decision network for wing expansion in Drosophila using targeted expression of the TRPM8 channel, J. Neurosci. 29, 3343–3353; DOI: 10.1523/JNEUROSCI.4241-08.2009.

60. Petri J, Syed MH, Rey S & Klämbt C (2019). Non-Cell-Autonomous Function of the GPI-Anchored Protein Undicht during Septate Junction Assembly, Cell Rep 26, 1641–1653.e4; DOI: 10.1016/j.celrep.2019.01.046.

61. Pircs K, Nagy P, Varga A, Venkei Z, Erdi B, Hegedus K & Juhasz G (2012). Advantages and limitations of different p62-based assays for estimating autophagic activity in Drosophila, PLoS ONE 7, e44214; DOI: 10.1371/journal.pone.0044214.

62. Rahman MM, Islam MR, Yamin M, Islam MM, Sarker MT, Meem AFK, Akter A, Emran TB, Cavalu S & Sharma R (2022). Emerging Role of Neuron-Glia in Neurological Disorders: At a Glance, Oxid Med Cell Longev 2022, 3201644; DOI: 10.1155/2022/3201644.

63. Schaaf ZA, Tat L, Cannizzaro N, Green R, Rülicke T, Hippenmeyer S & Zarbalis KS (2022). WDFY3 mutation alters laminar position and morphology of cortical neurons, Mol Autism 13, 27; DOI: 10.1186/s13229-022-00508-3.

64. Schneider CA, Rasband WS & Eliceiri KW (2012). NIH Image to ImageJ: 25 years of image analysis, Nat Methods 9, 671–675; DOI: 10.1038/nmeth.2089.

65. Schulte J, Tepass U & Auld VJ (2003). Gliotactin, a novel marker of tricellular junctions, is necessary for septate junction development in Drosophila, J Cell Biol 161, 991–1000; DOI: 10.1083/jcb.200303192.

66. Schumann I & Triphan T (2020). The PEDtracker: An Automatic Staging Approach for Drosophila melanogaster Larvae, Front. Behav. Neurosci. 14, 612313; DOI: 10.3389/fnbeh.2020.612313.

67. Sepp KJ & Auld VJ (1999). Conversion of lacZ enhancer trap lines to GAL4 lines using targeted transposition in Drosophila melanogaster, Genetics 151, 1093–1101; DOI: 10.1093/genetics/151.3.1093.

68. Sepp KJ, Schulte J & Auld VJ (2001). Peripheral glia direct axon guidance across the CNS/PNS transition zone, Dev Biol 238, 47–63; DOI: 10.1006/dbio.2001.0411.

69. Sim J, Osborne KA, Argudo García I, Matysik AS & Kraut R (2019). The BEACH Domain Is Critical for Blue Cheese Function in a Spatial and Epistatic Autophagy Hierarchy, Front Cell Dev Biol 7, 129; DOI: 10.3389/fcell.2019.00129.

70. Simonsen A, Birkeland HCG, Gillooly DJ, Mizushima N, Kuma A, Yoshimori T, Slagsvold T, Brech A & Stenmark H (2004). Alfy, a novel FYVE-domain-containing protein associated with protein granules and autophagic membranes, J Cell Sci 117, 4239–4251; DOI: 10.1242/jcs.01287.

71. Skeath JB, Wilson BA, Romero SE, Snee MJ, Zhu Y & Lacin H (2017). The extracellular metalloprotease AdamTS-A anchors neural lineages in place within and preserves the architecture of the central nervous system, Development 144, 3102–3113; DOI: 10.1242/dev.145854.

72. Søreng K, Pankiv S, Bergsmark C, Haugsten EM, Dahl AK, La Ballina LR de, Yamamoto A, Lystad AH & Simonsen A (2022). ALFY localizes to early endosomes and cellular protrusions to facilitate directional cell migration, J Cell Sci 135; DOI: 10.1242/jcs.259138.

73. Stessman HAF, Xiong B, Coe BP, Wang T, Hoekzema K, Fenckova M, Kvarnung M, Gerdts J, Trinh S & Cosemans N, et al. (2017). Targeted sequencing identifies 91 neurodevelopmental-disorder risk genes with autism and developmental-disability biases, Nat Genet 49, 515–526; DOI: 10.1038/ng.3792.

74. Stewart BA, Atwood HL, Renger JJ, Wang J & Wu CF (1994). Improved stability of Drosophila larval neuromuscular preparations in haemolymph-like physiological solutions, J Comp Physiol A 175, 179–191; DOI: 10.1007/BF00215114.

75. Stork T, Engelen D, Krudewig A, Silies M, Bainton RJ & Klämbt C (2008). Organization and function of the blood-brain barrier in Drosophila, J. Neurosci. 28, 587–597; DOI: 10.1523/JNEUROSCI.4367-07.2008.

76. Szabó Á, Vincze V, Chhatre AS, Jipa A, Bognár S, Varga KE, Banik P, Harmatos-Ürmösi A, Neukomm LJ & Juhász G (2023). LC3-associated phagocytosis promotes glial degradation of axon debris after injury in Drosophila models, Nat Commun 14, 3077; DOI: 10.1038/s41467-023-38755-4.

77. Tiklová K, Senti K-A, Wang S, Gräslund A & Samakovlis C (2010). Epithelial septate junction assembly relies on melanotransferrin iron binding and endocytosis in Drosophila, Nat Cell Biol 12, 1071–1077; DOI: 10.1038/ncb2111.

78. Tu H-Y, Yuan B-S, Hou X-O, Zhang X-J, Pei C-S, Ma Y-T, Yang Y-P, Fan Y, Qin Z-H & Liu C-F, et al. (2021). α-synuclein suppresses microglial autophagy and promotes neurodegeneration in a mouse model of Parkinson’s disease, Aging Cell 20, e13522; DOI: 10.1111/acel.13522.

79. Vanden Broeck L, Naval-Sánchez M, Adachi Y, Diaper D, Dourlen P, Chapuis J, Kleinberger G, Gistelinck M, Van Broeckhoven C & Lambert J-C, et al. (2013). TDP-43 loss-of-function causes neuronal loss due to defective steroid receptor-mediated gene program switching in Drosophila, Cell Rep 3, 160–172; DOI: 10.1016/j.celrep.2012.12.014.

80. Wang C, Liu Z & Huang X (2012). Rab32 is important for autophagy and lipid storage in Drosophila, PLoS ONE 7, e32086; DOI: 10.1371/journal.pone.0032086.

81. Wang T, Guo H, Xiong B, Stessman HAF, Wu H, Coe BP, Turner TN, Liu Y, Zhao W & Hoekzema K, et al. (2016). De novo genic mutations among a Chinese autism spectrum disorder cohort, Nat Commun 7, 13316; DOI: 10.1038/ncomms13316.

82. Winkler B, Funke D, Benmimoun B, Spéder P, Rey S, Logan MA & Klämbt C (2021). Brain inflammation triggers macrophage invasion across the blood-brain barrier in Drosophila during pupal stages, Sci Adv 7, eabh0050; DOI: 10.1126/sciadv.abh0050.

83. Wu X & Tu BP (2011). Selective regulation of autophagy by the Iml1-Npr2-Npr3 complex in the absence of nitrogen starvation, Mol Biol Cell 22, 4124–4133; DOI: 10.1091/mbc.E11-06-0525.

84. Xiong WC, Okano H, Patel NH, Blendy JA & Montell C (1994). repo encodes a glial-specific homeo domain protein required in the Drosophila nervous system, Genes Dev 8, 981–994; DOI: 10.1101/gad.8.8.981.

85. Yang M, Wang H, Chen C, Zhang S, Wang M, Senapati B, Li S, Yi S, Wang L & Zhang M, et al. (2021). Glia-derived temporal signals orchestrate neurogenesis in the Drosophila mushroom body, Proc Natl Acad Sci U S A 118; DOI: 10.1073/pnas.2020098118.

