## Supplemental Figures for "*Drosophila Bchs* overexpression recapitulates human *WDFY3* neurodevelopmental phenotypes with implications for glial cell involvement in altered head circumference"

### Supplemental Information

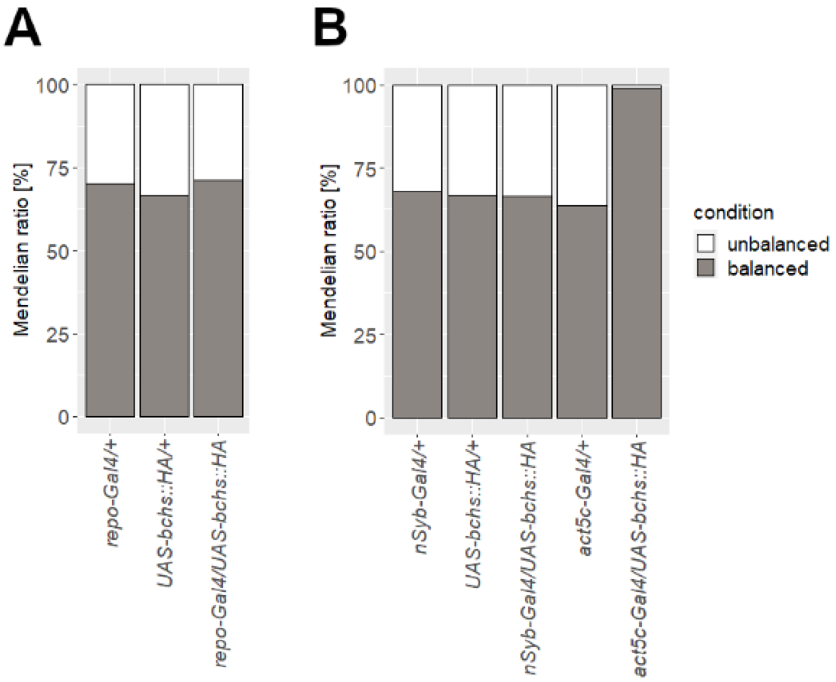


**Fig S1. Ubiquitous Bchs overexpression was lethal.**

A, B) Proportion of unbalanced (white) and balanced (gray) offspring. Unbalanced flies carried both, the *Gal4*-driver and the *UAS*-target gene, therefore, they overexpressed *Bchs* in the respective tissue. Balanced flies carried only one of both, *Gal4* or *UAS*, meaning that they did not overexpress *Bchs*. In neurons (B) or glial cells (A), *Bchs* overexpression did not affect the Mendelian ratio. In contrast, almost no adult flies existed that overexpressed *Bchs* ubiquitously (~1 %) (B), indicating the lethality of ubiquitous *Bchs* overexpression. Number of assessed individuals for each genotype ranged between 248−567 flies.

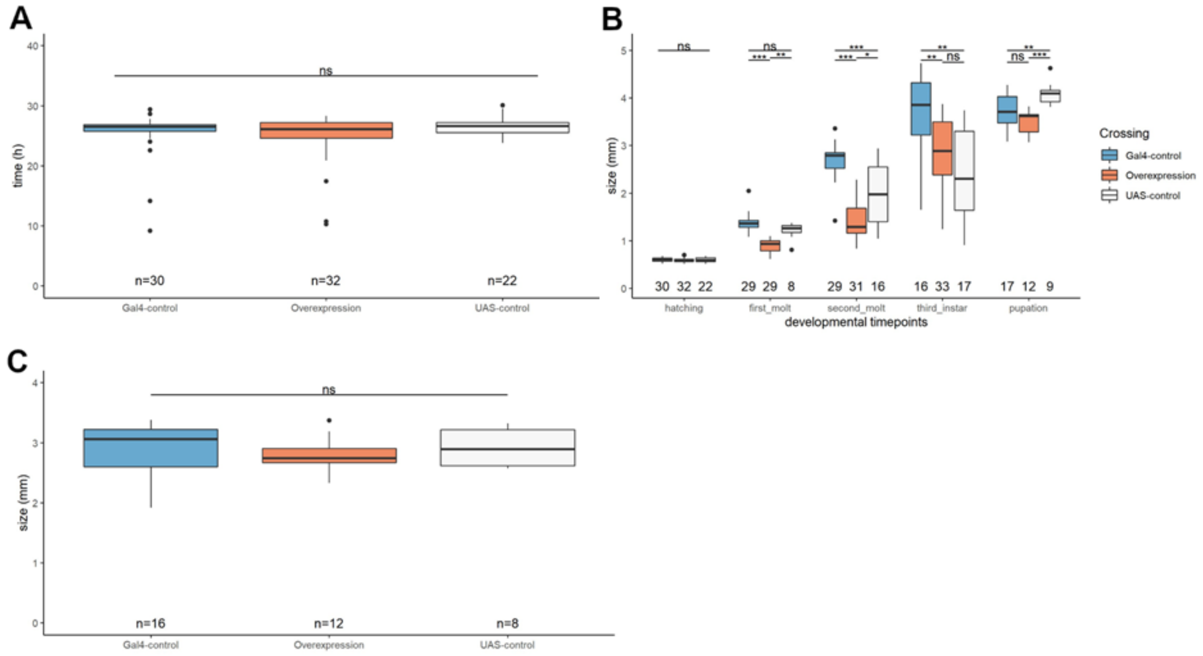


**Fig S2. Neuronal Bchs overexpression reduced larval size in early larval stages.**

A) Duration from egg laying to hatching of larvae. Neuronal *Bchs* overexpression larvae take a similar duration from egg laying to larval hatching as control genotypes. (Kruskal-Wallis test: $p$ = 0.51, $\text{mean}_{\text{Overexpression}}$ = 24.83 h, $\text{mean}_{\textit{Gal4}\text{-control}}$ = 25.36 h, $\text{mean}_{\textit{UAS}\text{-control}}$ = 26.62 h). B) Larval size was measured at different timepoints. Numbers at the bottom display the number of individual larvae measured. Neuronal *Bchs* overexpression larvae displayed similar lengths as controls at timepoint of hatching (One-way ANOVA: $p$ = 0.625). Larvae were significantly smaller at the timepoint of the first (62.44 h) (Mann Whitney U test, $p_{\text{overexpression}/\textit{Gal4}\text{-control}}$ < 0.001, $p_{\text{overexpression}/\textit{UAS}\text{-control}}$ = 0.002, $p_{\textit{Gal4}\text{-}/\textit{UAS}\text{-control}}$ = 0.12) and second (100.8 h) (Mann Whitney U test, $p_{\text{overexpression}/\textit{Gal4}\text{-control}}$ < 0.001, $p_{\text{overexpression}/\textit{UAS}\text{-control}}$ = 0.015, $p_{\textit{Gal4}\text{-}/\textit{UAS}\text{-control}}$ < 0.001) molt, but not at later stages (153 h) (Mann Whitney U test, $p_{\text{overexpression}/\textit{Gal4}\text{-control}}$ = 0.006, $p_{\text{overexpression}/\textit{UAS}\text{-control}}$ = 0.413, $p_{\textit{Gal4}\text{-}/\textit{UAS}\text{-control}}$ = 0.003) (and shortly before pupation, t-test, $p_{\text{overexpression}/\textit{Gal4}\text{-control}}$ = 0.363, $p_{\text{overexpression}/\textit{UAS}\text{-control}}$ < 0.001, $p_{\textit{Gal4}\text{-}/\textit{UAS}\text{-control}}$ = 0.007). C) Neuronal *Bchs* overexpression pupae had similar lengths as controls (Kruskal-Wallis test: $p$ = 0.6). A, B, C) Overexpression: *nSyb-Gal4/UAS-bchs::HA* (middle, orange), *Gal4*-control: *nSyb-Gal4/+* (left, blue), *UAS*-control: *UAS-bchs::HA/+* (right, white).

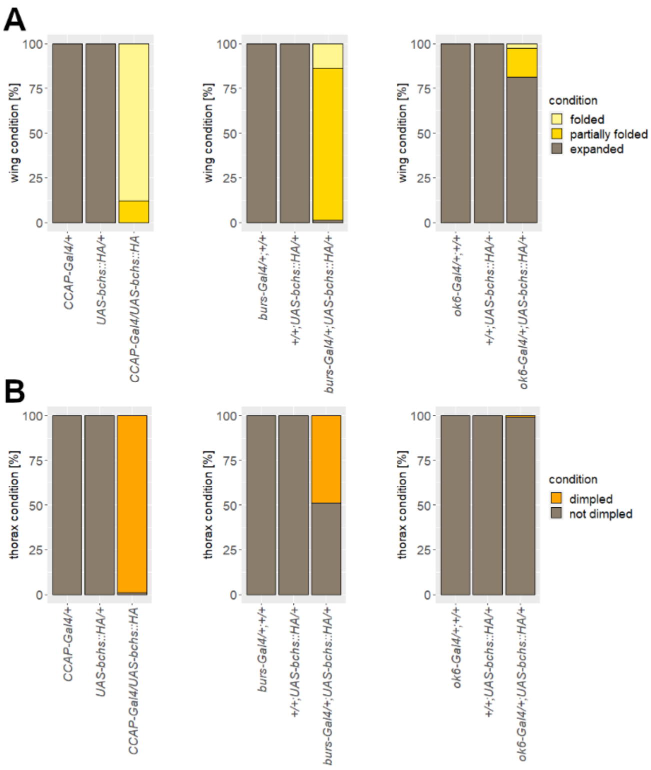


**Fig S3. Bchs overexpression in CCAP neurons caused wing and thorax abnormalities.**

A, B) *Bchs* overexpression caused wing (A) and thorax (B) defects. Overexpressing *Bchs* in the subset of CCAP neurons (left) resulted in a higher proportion of flies affected than *Bchs* overexpression in Burs neurons (middle) or motoneurons (*ok6-Gal4*, right). Number of assessed individuals for each genotype ranged between 144−412 flies.

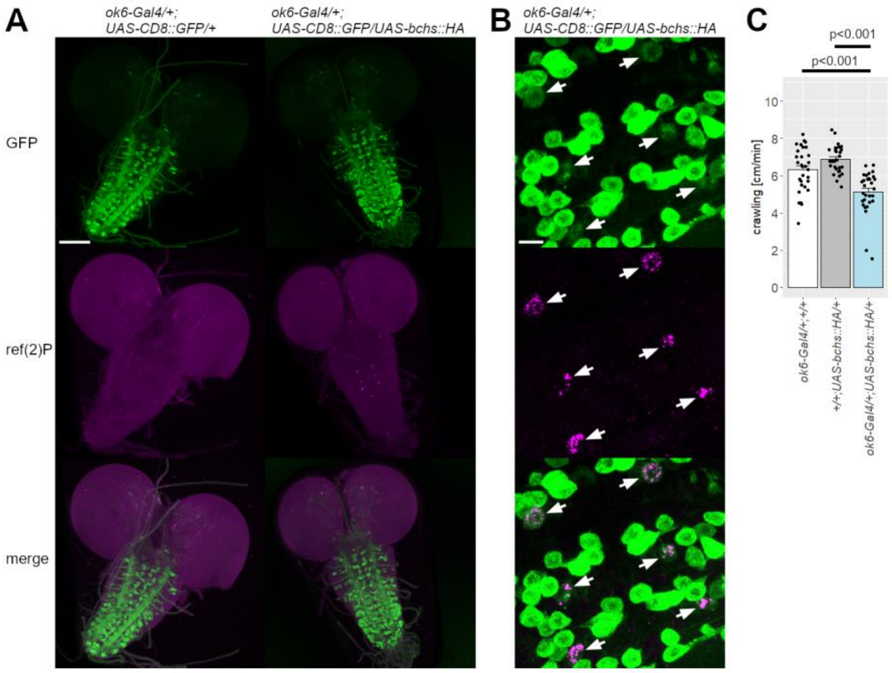


**Fig S4. Bchs overexpression in motoneurons impaired autophagy and larval locomotion.**

A, B) Membrane-bound GFP (green) was expressed in motoneurons and larval brains stained against ref(2)P (magenta). *Bchs* overexpression in motoneurons caused ref(2)P accumulation in a subset of neurons. The colocalization of GFP and ref(2)P signals (B, arrows) demonstrates that the ref(2)P-positive neurons are a subset of motoneurons. A) Scale bar: 100 µm. B) Insets were digitally multiplied to increase low intensity signals. Scale bar: 10 µm. C) *Bchs* overexpression in motoneurons (light blue) decreased the crawled distance of larvae in a given time. n = 30. Data are shown as mean ± SEM.

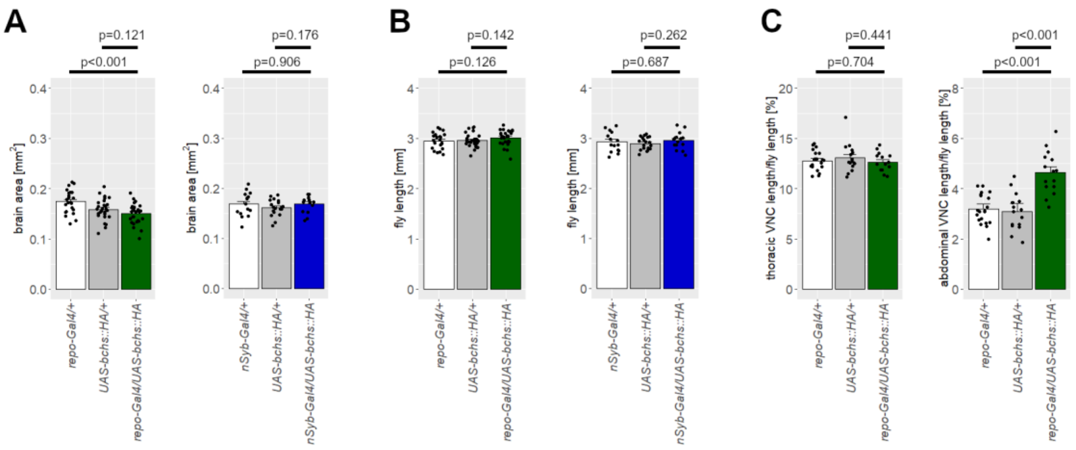


**Fig S5. CNS size and fly length.**

A) The size of the adult brain was not significantly altered by *Bchs* overexpression in glial cells (left, green) or neurons (right, blue) compared to controls. B) The length of the adult fly was not affected by *Bchs* overexpression in glial cells (left) or neurons (right). A, B) Glial dataset: n ≥ 23. Neuronal dataset: n ≥ 15. C) In adult flies overexpressing *Bchs* in glial cells, the length of the thoracic neuromeres of the VNC was not elongated (left). However, the length of the abdominal neuromeres was increased (right). n ≥ 15. Data are shown as mean ± SEM.

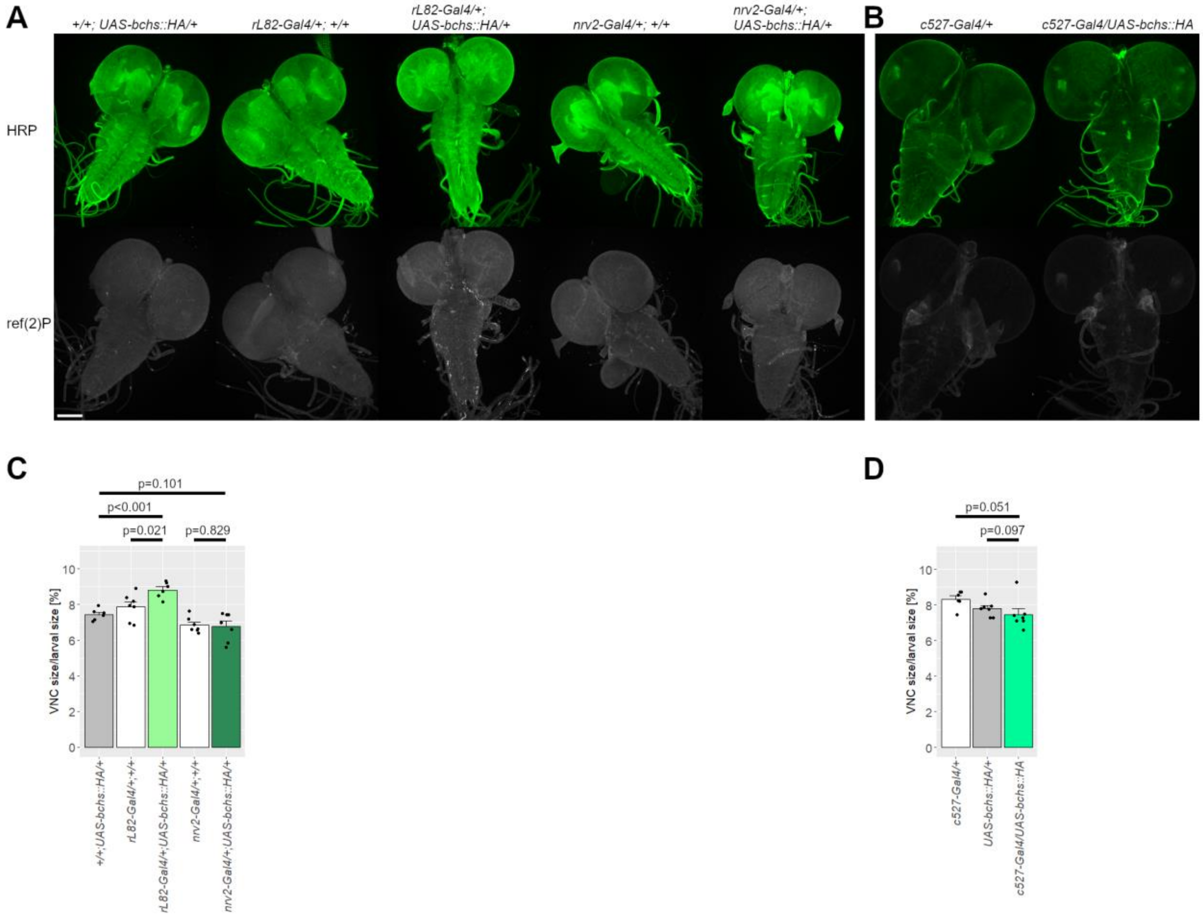


**Fig S6. Bchs overexpression in subperineural glia elongated the larval VNC.**

A, B) Larval brains were stained against HRP and ref(2)P. Scale bar: 100 µm. C, D) The VNC length of larval CNS overexpressing *Bchs* in subperineural (*rL82-Gal4*, C), wrapping (*nrv2-Gal4*, C) or perineural glia (*c527-Gal4*, D) was measured. An increase in VNC length was observed in larvae which overexpressed *Bchs* in subperineural glia. n > 6. Data are shown as mean ± SEM.

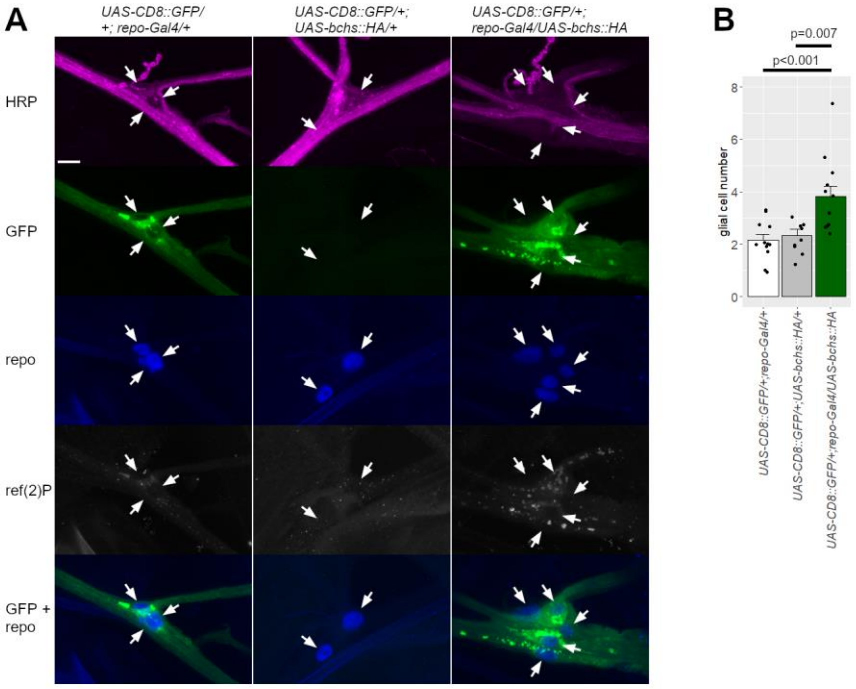


**Fig S7. Glial Bchs overexpression increased the number of glial cells in the peripheral nervous system.**

A) Larvae expressing membrane-bound GFP (green) in glial cells were stained against HRP (magenta), repo (blue) and ref(2)P (white). A control which did not express GFP was included (middle, *UAS-CD8::GFP/+; UAS-bchs::HA/+*). Images were taken from the peripheral nerve bundle innervating hemisegment A4R or A4L. Glial nuclei are indicated by arrows. A merge of GFP and repo signals is displayed at the bottom. Scale bar: 10 µm. B) Quantification of repo-positive nuclei number at the nerve bundle branching region. Glial *Bchs* overexpression larvae presented with a gain in glial cell number. $n \geq 9$. Data are shown as mean ± SEM.

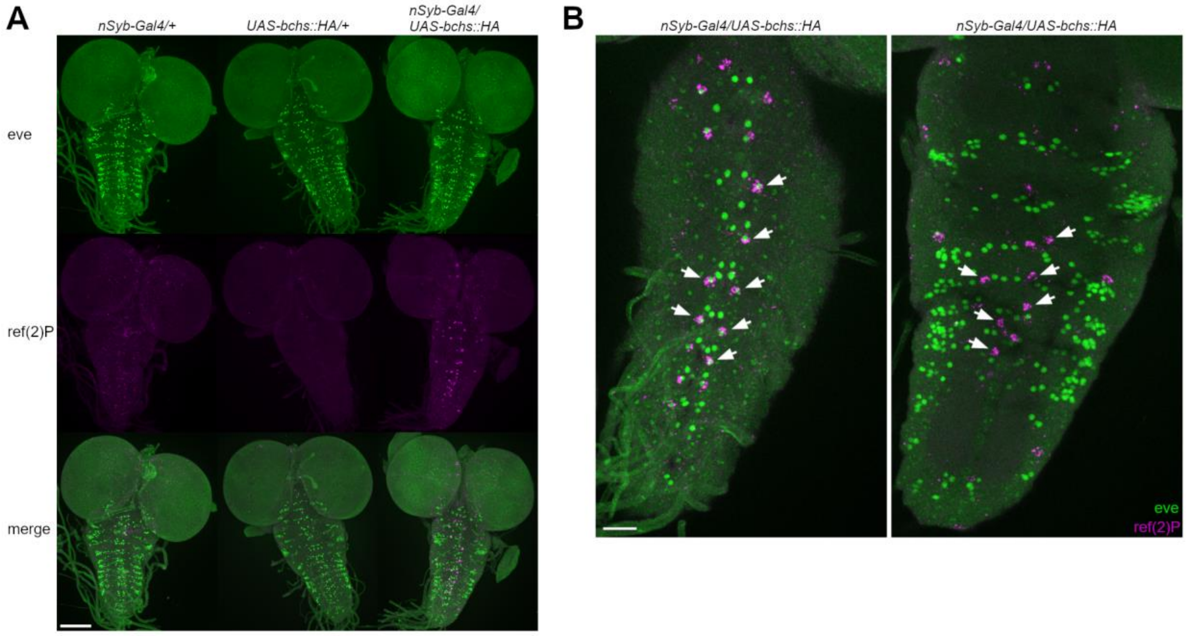


**Fig S8. Subset of eve neurons accumulated ref(2)P in response to Bchs overexpression.**

A, B) Larval brains were stained against eve (green) and ref(2)P (magenta). A) Scale bar: 100 µm. B) Insets to the VNC of neuronal *Bchs* overexpressing brains. Signals of eve and ref(2)P were merged to display colocalization. Images show a maximum projection of only a subset of confocal imaging layers. Some ref(2)P aggregates did localize to eve neurons (left, arrows), however, there was also ref(2)P accumulation that did not correspond to eve neurons (right, arrows). Therefore, ref(2)P aggregates caused by neuronal *Bchs* overexpression are located to a subset of eve neurons but also to another group of neurons. Scale bar: 25 µm.

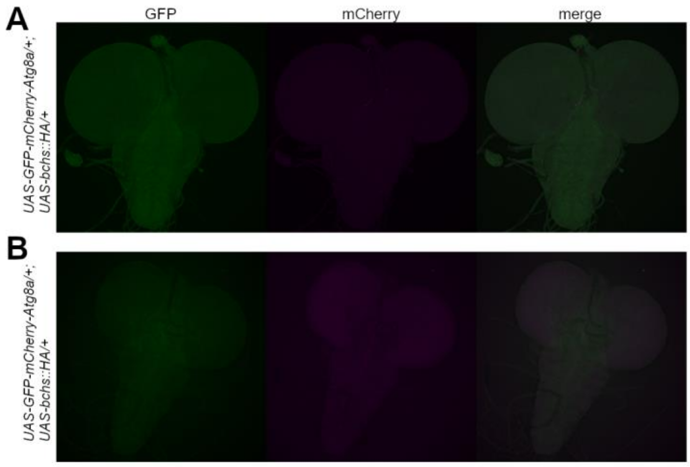


**Fig S9. Controls for GFP-mCherry-Atg8a reporter assay.**

A, B) Negative controls for GFP-mCherry-Atg8a reporter assays only carrying *UAS*-target genes but no *Gal4*-driver. A) glial *Bchs* overexpression dataset. B) neuronal *Bchs* overexpression dataset. Related to Figure 5.

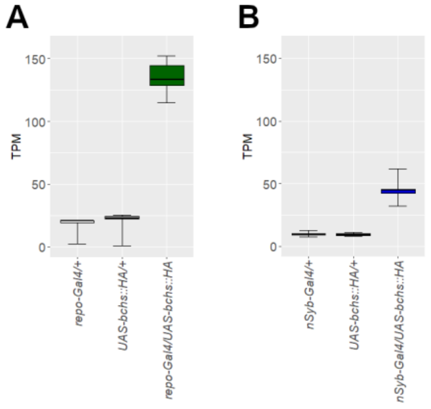


**Fig S10. Transcript levels of Bchs.**

A, B) Expression levels of *Bchs* transcripts in glial *Bchs* overexpression flies (A, green, *repo-Gal4/UAS-bchs::HA*), neuronal *Bchs* overexpression flies (B, blue, *nSyb-Gal4/UAS-bchs::HA*) and controls. white: *repo-Gal4/+* (A) or *nSyb-Gal4/+* (B), grey: *UAS-bchs::HA/+*. n = 5. TPM: transcripts per million. Horizontal line represents the median. Lower and upper hinges correspond to the 25$^{th}$ and 75$^{th}$ percentiles. Whiskers show maximum and minimum values.

**Tab S1:**

Differentially expressed genes (DEGs) of *Bchs* overexpression flies and GO enrichment analysis from DEGs. Gene expression levels from glial and neuronal *Bchs* overexpression flies (*repo-Gal4/UAS-bchs::HA* and *nSyb-Gal4/UAS-bchs::HA*, respectively) was compared to the respective controls (glial: *repo-Gal4/*+ and *UAS-bchs::HA/*+, neuronal: *nSyb-Gal4/*+ and *UAS-bchs::HA/+)*. Genes were considered to be significantly differentially expressed if adjusted *p*-value < 0.05. GO modules were considered to be significantly enriched with a *p*-value after family wise error rate multiple correction < 0.05.
